# Transcriptional programs mediating neuronal toxicity and altered glial-neuronal signaling in a *Drosophila* knock-in tauopathy model

**DOI:** 10.1101/2024.02.02.578624

**Authors:** Hassan Bukhari, Vanitha Nithianandam, Rachel A. Battaglia, Anthony Cicalo, Souvarish Sarkar, Aram Comjean, Yanhui Hu, Matthew J. Leventhal, Xianjun Dong, Mel B. Feany

## Abstract

Missense mutations in the gene encoding the microtubule-associated protein tau cause autosomal dominant forms of frontotemporal dementia. Multiple models of frontotemporal dementia based on transgenic expression of human tau in experimental model organisms, including *Drosophila*, have been described. These models replicate key features of the human disease, but do not faithfully recreate the genetic context of the human disorder. Here we use CRISPR-Cas mediated gene editing to model frontotemporal dementia caused by the tau P301L mutation by creating the orthologous mutation, P251L, in the endogenous *Drosophila tau* gene. Flies heterozygous or homozygous for tau P251L display age-dependent neurodegeneration, metabolic defects and accumulate DNA damage in affected neurons. To understand the molecular events promoting neuronal dysfunction and death in knock-in flies we performed single-cell RNA sequencing on approximately 130,000 cells from brains of tau P251L mutant and control flies. We found that expression of disease-associated mutant tau altered gene expression cell autonomously in all neuronal cell types identified and non-cell autonomously in glial cells. Cell signaling pathways, including glial-neuronal signaling, were broadly dysregulated as were brain region and cell-type specific protein interaction networks and gene regulatory programs. In summary, we present here a genetic model of tauopathy, which faithfully recapitulates the genetic context and phenotypic features of the human disease and use the results of comprehensive single cell sequencing analysis to outline pathways of neurotoxicity and highlight the role of non-cell autonomous changes in glia.

## Introduction

The neuronal microtubule-associated protein tau forms insoluble deposits termed neurofibrillary tangles and neuritic threads in neuronal soma and processes in a diverse group of age-dependent neurodegenerative diseases, including Alzheimer’s disease and frontotemporal dementia. These disorders have collectively been termed “tauopathies” (Feany and Dickson 1996; Götz et al. 2019; Goedert 2004). While wild type tau is deposited in Alzheimer’s disease and other more common tauopathies, missense mutations in tau occur in rarer familial forms of tauopathy causing neurodegeneration and insoluble tau deposition. Autosomal dominant disease-causing mutations occur throughout the tau protein but are particularly frequent in exon 10, which contains one of four microtubule binding repeats (Ghetti et al. 2015). These repeats mediate microtubule (Lee et al. 1989; Butner and Kirschner 1991) and actin (Cabrales Fontela et al. 2017) binding, and are important determinants of tau aggregation (von Bergen et al. 2000). Experimental models of tauopathy have been created in diverse model organisms, from yeast to non-human primates, by expressing wild type or frontotemporal dementia-associated mutant forms of human tau in transgenic animals. Mutant forms of tau are typically more toxic than wild type tau in transgenic model organisms. Work in these models has implicated a number of cellular pathways in mediating tau neurotoxicity, including mitochondrial dysfunction (Rhein et al. 2009; DuBoff et al. 2012), oxidative stress (Dias-Santagata et al. 2007; Dumont et al. 2011) and aberrant cell cycle reentry of postmitotic neurons (Khurana et al. 2006; Andorfer et al. 2005).

However, while transgenic models have been useful, they do not faithfully replicate the genetic underpinnings of the authentic human disorders and thus may not allow the identification and study of the full complement of important mediators of tauopathy pathogenesis. We have therefore used CRISPR-Cas9 gene editing to model familial frontotemporal dementia caused by missense mutations in tau more precisely in *Drosophila*. Mutation of proline 301 to leucine in exon 10 is the most common mutation of tau in frontotemporal dementia patients (Poorkaj et al. 2001), and has been frequently modeled in transgenic animals (Goedert and Jakes 2005). The overall structure and expression of tau is conserved from mammals to *Drosophila* (Heidary and Fortini 2001), with proline 251 being orthologous to human proline 301. We have therefore replaced *Drosophila* tau proline 251 with leucine (P251L) and phenotypically analyzed the resultant homozygous and heterozygous animals with age. We have additionally performed single-cell sequencing to identify cell populations, networks and signaling systems altered by mutant tau expression.

## Results

### Phenotypic analysis of a *Drosophila* knock-in model of frontotemporal dementia

We used CRISPR-Cas9 gene editing to recapitulate the genetic basis of human frontotemporal dementia in the powerful genetic experimental organism *Drosophila* by modeling the disease-causing proline 301 to leucine in fly tau. Protein sequence alignment shows that the microtubule-binding domains, including human tau proline 301 are evolutionary conserved from *Drosophila* to humans (Supplemental Fig. S1). The homologous residue of the human tau proline 301, *Drosophila* tau proline 251, was mutated to leucine using a highly efficient guide RNA along with single-stranded oligodeoxynucleotides (Fig. 1A,B). Mutant tau was expressed at equivalent levels to wild type tau (Supplemental Fig. 2A).

**Figure 1.**
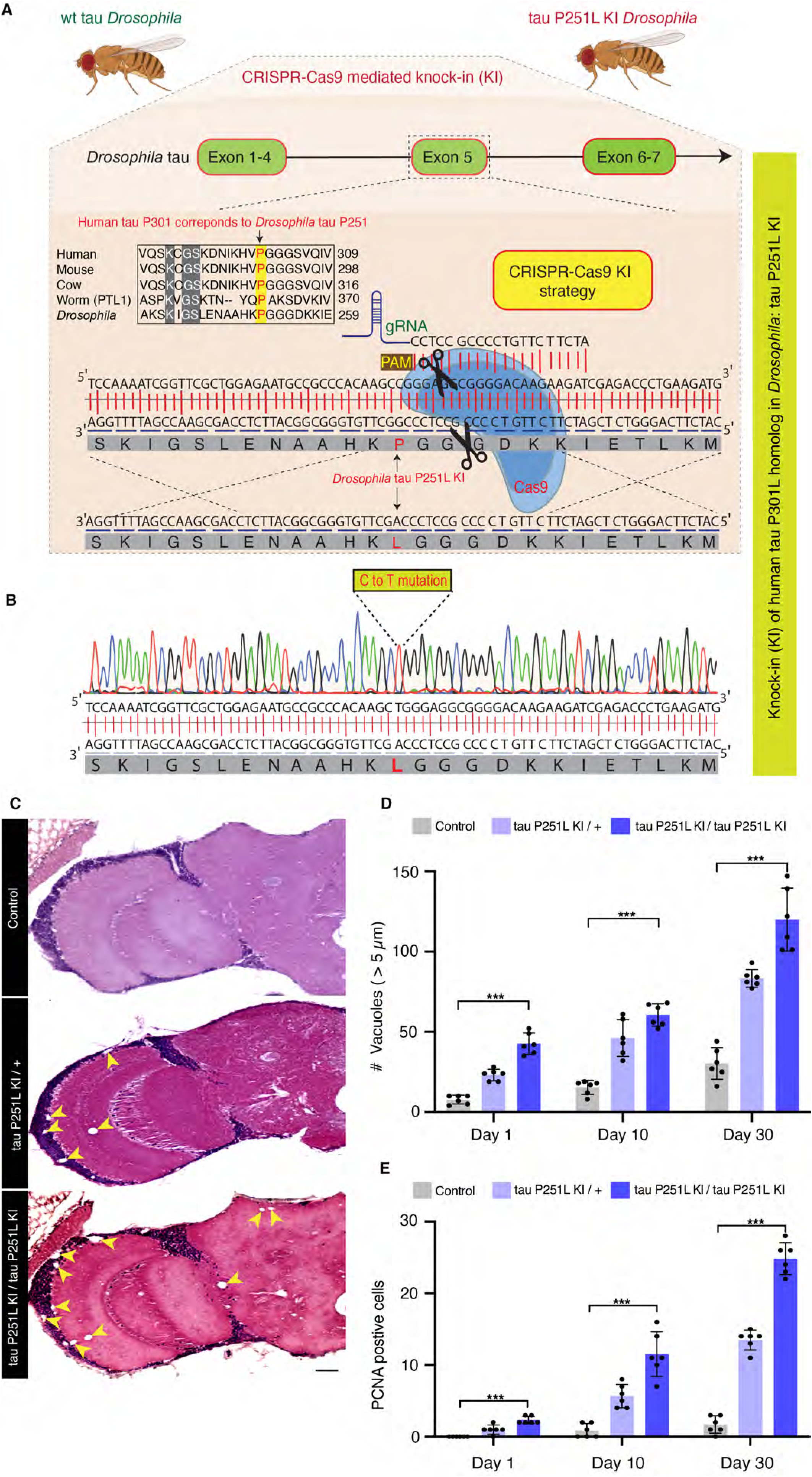
CRISPR-Cas9-mediated knock-in model of frontotemporal dementia in *Drosophila*. CRISPR-Cas9 gene editing strategy to knock in the human tau P301L homologous mutation in *Drosophila,* tau P251L, located in exon 5 of *Drosophila* tau (*A*). Successful mutation in homozygous tau P251L knock-in flies (*B*). Hematoxylin and eosin staining reveals evidence of neurodegeneration as seen by increased number of brain vacuoles (arrowheads) with age in homozygous and heterozygous knock-in animals (*C,D*). Scale bar represents 10 µm (*C*). Neurodegeneration is accompanied by abnormal cell cycle reentry as marked by proliferating cell nuclear antigen (PCNA) staining (*E*). Flies are 30 days old in (*C*) and the age indicated in the figure labels in (*D,E*). n = 6 per genotype and time point (*D,E*). Data are presented as mean ± SD. *** = P < 0.001, one-way ANOVA with Tukey post-hoc analysis.

Expression of frontotemporal dementia-linked forms of mutant tau, including P301L, lead to age-dependent neuronal loss in patients and in transgenic models (Ghetti et al. 2015; Lewis et al. 2001; Yoshiyama et al. 2007; Götz et al. 2001). We thus examined the histology of brains of heterozygous (P251L / +) and homozygous (P251L) tau knock-in animals with age. We found increased numbers of cortical and neuropil vacuoles in knock-in animals (Fig. 1C,D). Neurodegeneration in *Drosophila* is frequently accompanied by the formation of brain vacuoles (Buchanan and Benzer 1993; Wittmann et al. 2001; Ordonez et al. 2018; Heisenberg and Böhl 1979). Increasing numbers of vacuoles were observed with advancing age, and with two copies of the P251L compared with one copy (Fig. 1C,D). Inappropriate neuronal cycle reentry is a feature of human tauopathy (Husseman et al. 2000) and human tau transgenic animals (Andorfer et al. 2005; Khurana et al. 2006). We stained control and tau P251L knock-in brains with an antibody directed to proliferating cell nuclear antigen (PCNA) to assess cell cycle activation (Khurana et al. 2006). We observed increasing cell cycle reentry with age in tau P251L knock-in brains, with more cell cycle activation in homozygotes compared to heterozygotes (Fig. 1E, Supplemental Fig. 2B).

Metabolic alterations and mitochondrial dysfunction are pervasive features of neurodegenerative diseases, including tauopathies (DuBoff et al. 2013; Götz et al. 2019). We thus performed metabolic analysis on intact whole fly brains using the Seahorse XFe96 Analyzer (Neville et al. 2018). We observed reduced basal oxygen consumption rate (OCR) and a shift in mitochondrial bioenergetics to quiescent metabolic state in tau 251L knock-in animals, with homozygotes showing more impairment than heterozygotes (Fig. 2A,B).

**Figure 2.**
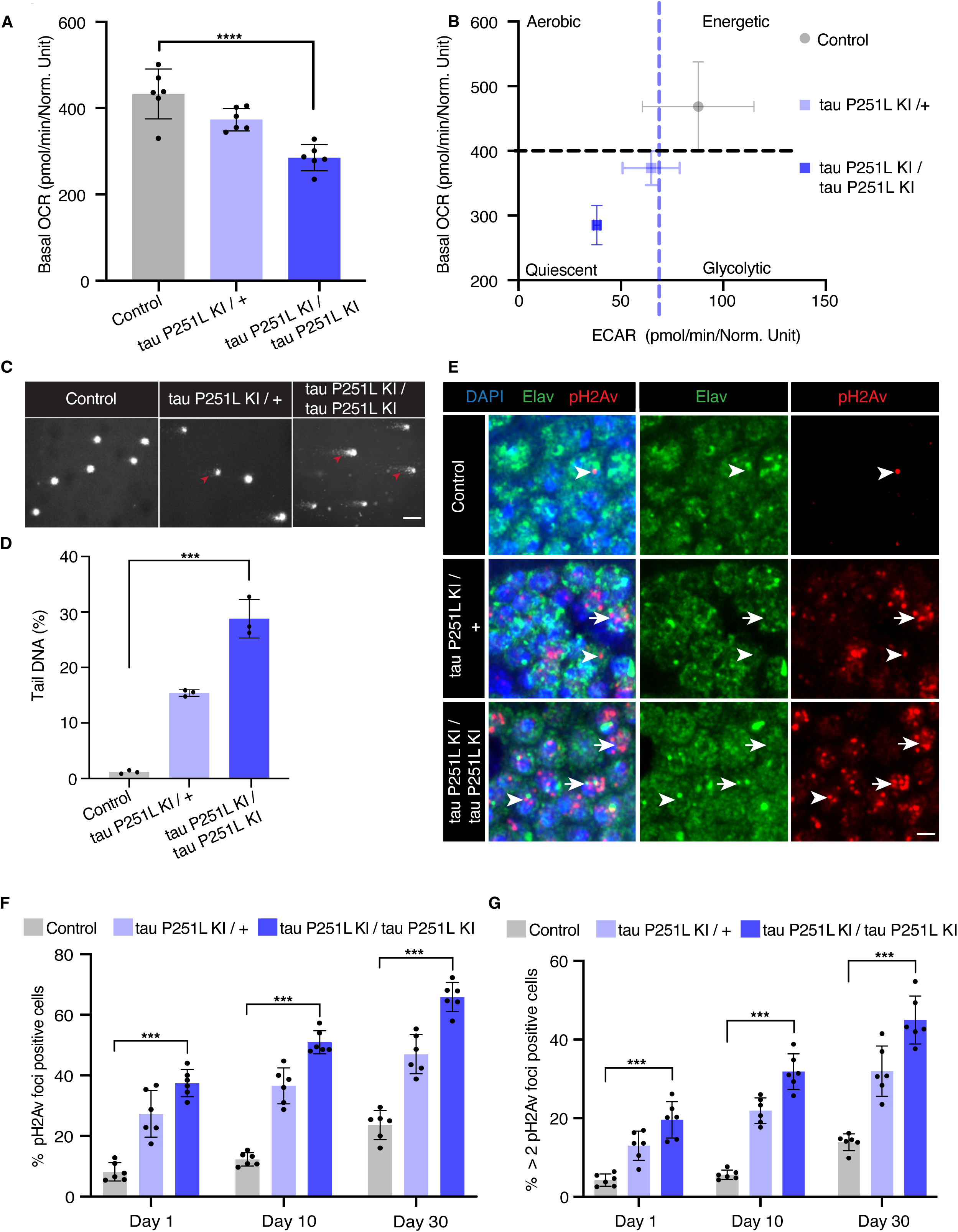
Mitochondrial dysfunction and DNA damage in tau P251L knock-in brains. Decreased oxygen consumption rate (OCR) (*A*) and shift to a quiescent metabolic phenotype as indicated by plotting the OCR vs. the extracellular acidification rate (ECAR) (*B*) in homozygous and heterozygous tau P251L knock-in brains compared to controls. n = 6 per genotype. Elevated levels of DNA damage in tau P251L knock-in brains as indicated by increased tail length (*C*, arrowheads, *D*) following electrophoresis of nuclei from dissociated brains in the comet assay. n = 3 per genotype. Increase in the number of Kenyon cells neurons (*E*, identified by the neuronal marker elav) containing DNA double-strand breaks as marked by pH2Av foci (*E*, arrowheads; arrows indicate neuronal nuclei with more than two foci) in histological sections of mushroom bodies (Kenyon cells) from tau P251L knock-in animals, as quantified in (*F,G*). n = 6 per genotype and time point. Scale bars represent 5 µm. Flies are 10 days old in (*A-D*), 30 days old in (*E*) and the age indicated in the figure labels in (*F,G*). Data are presented as mean ± SD. *** = P < 0.001, one-way ANOVA with Tukey post-hoc analysis.

Oxidative stress accompanying mitochondrial dysfunction results in damage to key cellular substrates, including DNA. DNA damage commonly occurs in age-related neurodegenerative diseases (Welch and Tsai 2022), including tauopathies (Khurana et al. 2012; Thadathil et al. 2021; Shanbhag et al. 2019). We took two approaches to examining DNA damage in tau P251L knock-in animals. First, we used the comet assay, in which DNA single- or double-strand breaks are demonstrated using single-cell gel electrophoresis (Khurana et al. 2012; Frost et al. 2014). We observed that nuclei from brains of tau P251L knock-in flies displayed almost 2-fold longer comet tails than controls (Fig. 2C, arrowheads, D).

As a second measure of DNA damage, we immunostained for the histone variant H2Av phosphorylated at serine 137 (pH2Av), a marker of DNA double-strand breaks (Madigan et al. 2002; Khurana et al. 2012; Frost et al. 2014). We found significantly increased numbers of double-strand breaks within neurons (Fig. 2E, arrows, arrowheads, F,G). DNA double-strand breaks were elevated with age and in homozygous compared to heterozygous tau P251L knock-in flies (Fig. 2E-G). Increased DNA damage was assessed by counting both the numbers of Kenyon cell nuclei containing pH2Av foci, and the number of Kenyon cell nuclei containing more than 2 foci (Fig. 2E, arrows, G), which correlates with increased numbers of DNA double-strand breaks (Lapytsko et al. 2015; Hong and Choi 2013).

### Single-cell RNA sequencing reveals gene expression changes mediated by pathologic tau

Our tau P251L knock-in flies replicate important features of human tauopathies and transgenic models of the disorders. We therefore performed single-cell RNA sequencing to investigate transcriptional programs and cellular pathways altered by expression of mutant tau. Using an optimized brain dissociation method, 10x library preparation, sequencing, and a bioinformatics analysis pipeline, we implemented single-cell RNA sequencing on tau P251L knock-in and control *Drosophila* brains at 10 days of age (Fig. 3A). The 10-day time point was chosen to identify early perturbations related to neuronal dysfunction and degeneration (Fig. 1,2). After stringent quality control 130,489 high-quality cells were retained in the final integrated dataset and 29 clusters of cells were identified. We annotated 26 clusters using a published fly cell atlas (Li et al. 2022). We used the most highly expressed marker genes within each cluster to identify clusters. For instance, we used *dac*, *crb* and *jdp* to annotate Kenyon cells; *Yp1*, *Yp2* and *Yp3* for mushroom body output neurons (MBON); *Mtna*, *CG8369* and *CG1552* for glia; *CG34355*, *Gad1* and *mamo* for medullary neurons; *acj6*, *Li1* and *sosie* for T neurons (Supplemental Figs. S3,S4,S5). The clustered dot plot illustrates enrichment of marker genes in annotated neuronal and glial clusters (Fig. 3C, Supplemental Fig. S4C). Based on prior published analyses (Li et al. 2022; Croset et al. 2018; Davie et al. 2018) we further outlined major groups of cells, including Kenyon cells, medullary neurons, mushroom body output neurons (MBON), astrocytes and perineurial glia (Fig. 3B; Supplemental Fig. S3; Supplemental Table S1). As previously observed (Davie et al. 2018), cholinergic neurons were the most common neuronal type defined by neurotransmitter phenotype, followed by GABAergic and glutamatergic neurons (Supplemental Fig. S3). Less abundant clusters of dopaminergic neurons were also identified (Supplemental Fig. S3). In summary, our scRNA sequencing in a precisely edited *Drosophila* tauopathy model, yielded 130,489 high quality cells, and identified cellular populations throughout diverse brain regions and cell types, including rarer cell populations such as astrocytes and perineurial glia.

**Figure 3.**
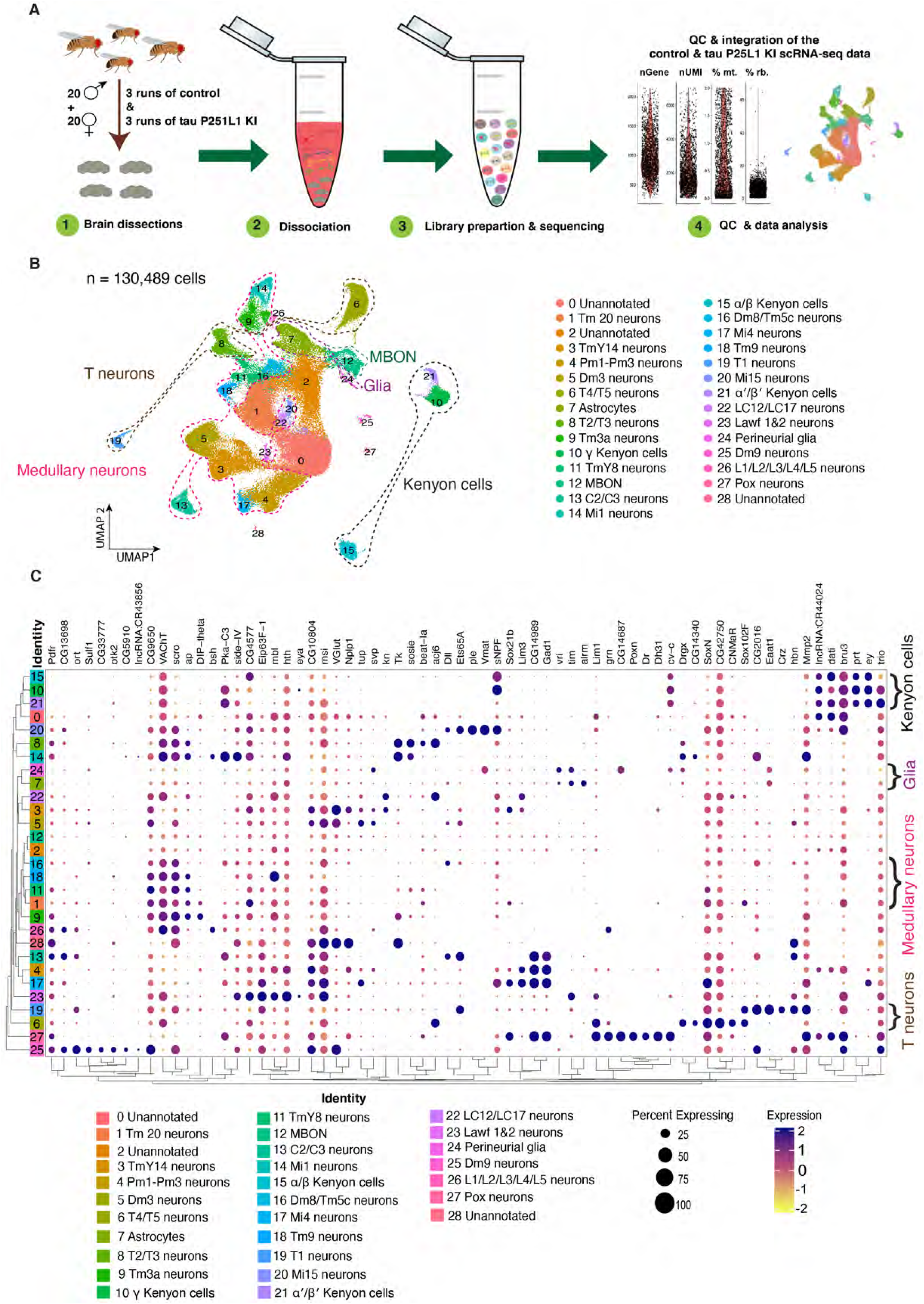
Single-cell RNA sequencing of tau P251L knock-in brains. Schematic of the single-cell RNA sequencing analysis pipeline (*A*). Following dissection, brains were dissociated in the enzymatic solutions and the single-cell suspension was encapsulated by 10x chromium platform. The 10x libraries were prepared, sequenced and after quality control, data was analyzed. UMAP representation of the 6 integrated sc-RNA sequencing runs, 3 control and 3 tau P251L knock-in (*B*). The integrated dataset contains 130,489 cells, and 26 clusters out of 29 were annotated. Percentage expression heatmap of the highly expressed marker genes within all clusters (*C*). Flies are 10 days old.

After sample integration, quality control and cluster annotation, we performed differential gene expression analysis (DEG) to identify genes modulated by precise pathologic mutation modeling of tauopathy in the *Drosophila* brain. DEG analysis of all the 26 annotated clusters revealed that tau P251L knock-in altered genes throughout the *Drosophila* brain, in both neurons and glia (Fig. 4A, Supplemental Table S2). We found that 472 genes were upregulated across all clusters in tau P251L knock-in brains, while 1145 genes were downregulated (Supplemental Table S3). Interestingly, transposable elements (*FBti0020120 RR48373-transposable-element, FBti0063007, FBti0019000, FBti0019150, RR50423-transposable-element, FBti0019148*) were frequently upregulated in P251L knock-in brains (Fig. 4B), consistent with findings from *Drosophila* human tau overexpression models and human Alzheimer’s disease brain tissue (Sun et al. 2018; Guo et al. 2018). The set of commonly downregulated genes was notable for multiple ribosomal protein genes (Fig. 4C), suggesting a translational defect in tauopathy. Multiple nuclear and mitochondrially encoded respiratory chain subunits and other mitochondrial proteins were notably present in the commonly upregulated and downregulated gene, as were genes encoding cytoskeletal and associated proteins (*Arc1, Msp300, Ank2, unc-104, Amph, brp, alphaTub84B*). Both categories of genes fit well with known mediators of tauopathy pathogenesis (DuBoff et al. 2013; Schulz et al. 2023).

**Figure 4.**
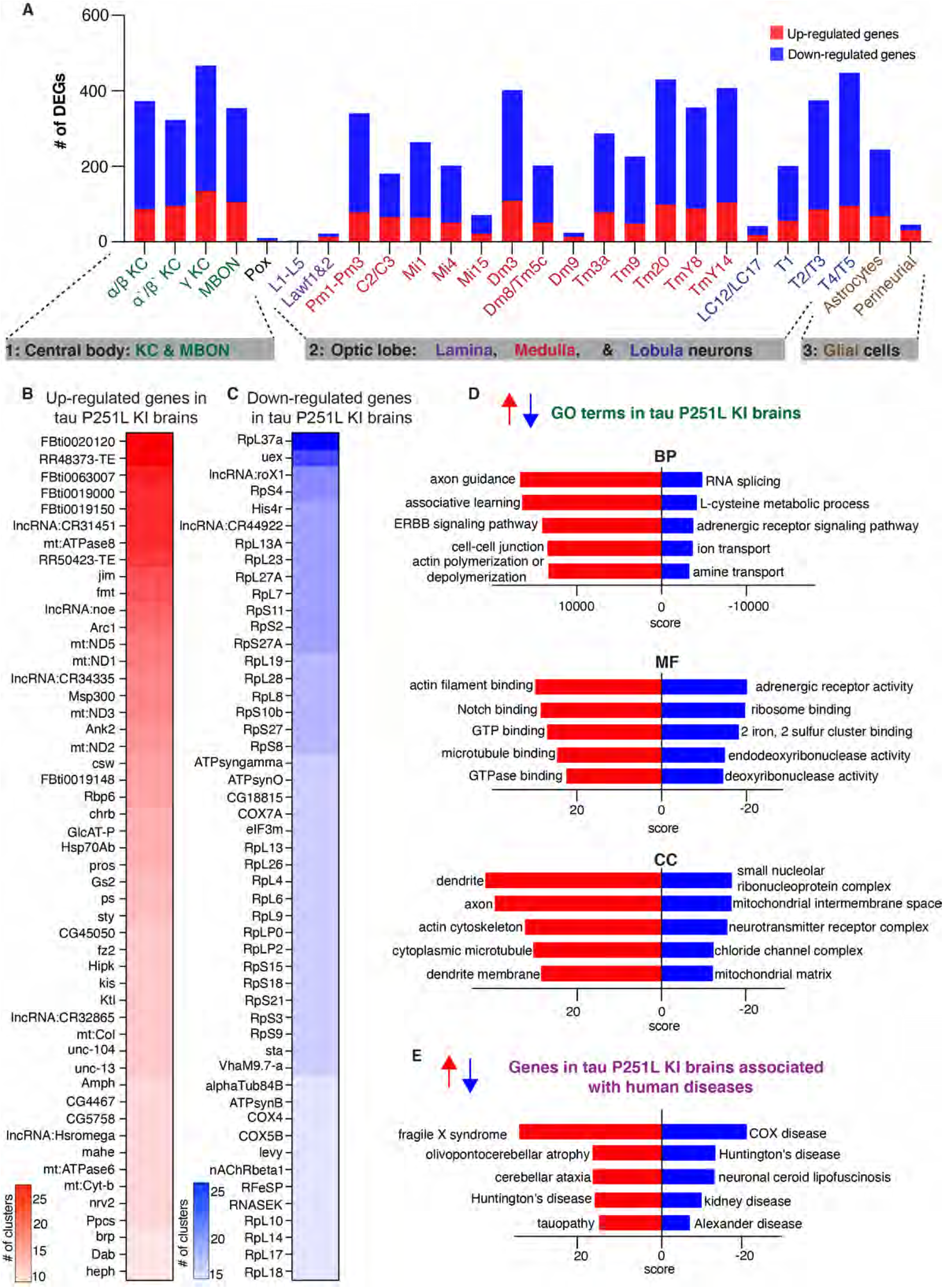
Differential gene expression and enrichment analysis of the scRNA seq dataset in tau P251L knock-in brains compared to controls. The number of differentially expressed genes (DEGs), both upregulated and downregulated genes, in all the annotated clusters of tau P251L knock-in brains compared to controls (*A*). Results are displayed across three major anatomic and functional classes of cells: 1) central body containing three clusters of Kenyon cells (KC), mushroom body output neurons and pox neurons, 2) optic lobe neurons containing lamina, medullary and lobula neurons clusters, and 3) glia cells containing astrocytes and perineurial clusters. Heatmap of the top 50 upregulated (*B*) and downregulated (*C*) genes in all the clusters of tau P251L knock-in brains compared to controls (Supplemental Table S3). Gene ontology (GO) enrichment analysis identified top upregulated and downregulated biological processes (BP), molecular functions (MF), and cellular components (CC) (*D*). Analysis of human disease associated genes revealed top upregulated and downregulated disease-associated gene sets (*E*). Score represents the combined score c = log(p)*z (Chen et al. 2013).

As expected, gene enrichment analyses (Fig. 4D; Supplemental Fig. S6) highlighted mitochondrial and cytoskeletal processes. In addition, diverse metabolic and neuronal function pathways, including associated learning, previously associated with Alzheimer’s disease and related tauopathies emerged from gene ontology (GO) enrichment analyses. Interestingly, enrichment analysis for human disease associated genes revealed predominantly neurodegenerative disorders, including tauopathy (Fig. 4E).

### Distinct and shared region- and cell-specific transcriptional programs in tau P251L knock-in brains

Significant anatomic and cell type selectivity characterizes human neurodegenerative diseases, including tauopathies. We therefore analyzed gene expression changes separately in anatomically and functionally related groups of cells including the central body (Kenyon cells, MBON and Pox neurons), optic lobe (lamina, medulla, and lobula neurons) and glia (astrocytes and perineurial glia). Volcano plots in Supplemental Fig. 7A, C and F present upregulated and downregulated genes in each group of cells. Interestingly, transposable element were top up-regulated genes in each of the three groups. GO enrichment analyses (Supplemental Fig. 7B,D) identified distinct biological processes altered by mutant tau expression in the central body compared to the optic lobe. Both associative learning and cAMP metabolic process were specifically identified in the central body, correlating with the importance of Kenyon cells in learning and memory in flies and with the central role for cAMP underlying learning and memory (Guven-Ozkan and Davis 2014; Feany and Quinn 1995). Heterochromatin organization and DNA repair, both processes strongly implicated in tauopathy pathogenesis (Fig. 2) (Khurana et al. 2012; Frost et al. 2014; Welch and Tsai 2022) emerged as enriched processes following analysis of the central body and optic lobe separately (Supplemental Fig. 7D). Direct comparison of differentially regulated genes in central body compared to optic lobe neurons revealed 239 commonly regulated genes and 562 distinct genes (Supplemental Fig. 7E). Consistent with analysis of the total transcriptome (Fig. 4), shared biological processes included downregulation of mitochondrial genes and upregulation of axon guidance-associated genes (Supplemental Fig. 6E; Supplemental Table S4).

Since tau is a predominantly neuronal gene (Heidary and Fortini 2001; Goedert 2004; Götz et al. 2019) the observed changes in neuronal transcriptomes plausibly reflect cell-autonomous effects of frontotemporal dementia associated mutant tau protein. Interestingly, our single-cell approach also revealed significant changes in gene expression in glial cells in tau P251L knock-in brains (Supplemental Fig. 7F,G). Expression of mutant tau may thus exert non-cell autonomous control on glial transcriptional programs. Metabolic processes (Supplemental Fig. 7G) were downregulated in glia in response to neuronal expression of mutant tau, consistent with the importance of glial metabolism in supporting a wide array of neuronal functions (Nedergaard and Verkhratsky 2012; Verkhratsky et al. 2012). Interestingly, the top two GO processes identified by analysis of upregulated glial genes were associative learning and regulation of neuronal remodeling, suggesting that coordinate changes in neurons and glia may lead to impairment of critical neuronal functions when mutant tau is expressed by neurons.

We next constructed protein interaction networks to explore further the biological pathways altered in tau P251L knock-in brains compared to controls. We used the solution of the prize-collecting Steiner forest algorithm (Tuncbag et al. 2013) to map differentially expressed genes onto a network of physical protein interactions using *Drosophila* interactome data. Networks constructed from the central body, optic lobe and glial cells were substantially distinct (Fig. 5), consistent with differential effects of mutant tau on different anatomic regions and cell types. The electron transport chain was identified in subnetworks from both the optic lobe and glia suggesting that mutant tau can influence mitochondrial function in both a cell-autonomous and non-cell autonomous fashion (Figs. 2,5). Regulation of nuclear function was commonly identified in both central body and optic lobe neurons, consistent with a strong influence of neuronally expressed tau on chromatin structure mediated through the Linker of Nucleoskeleton and Cytoskeleton (LINC) complex (Frost et al. 2014, 2016).

**Figure 5.**
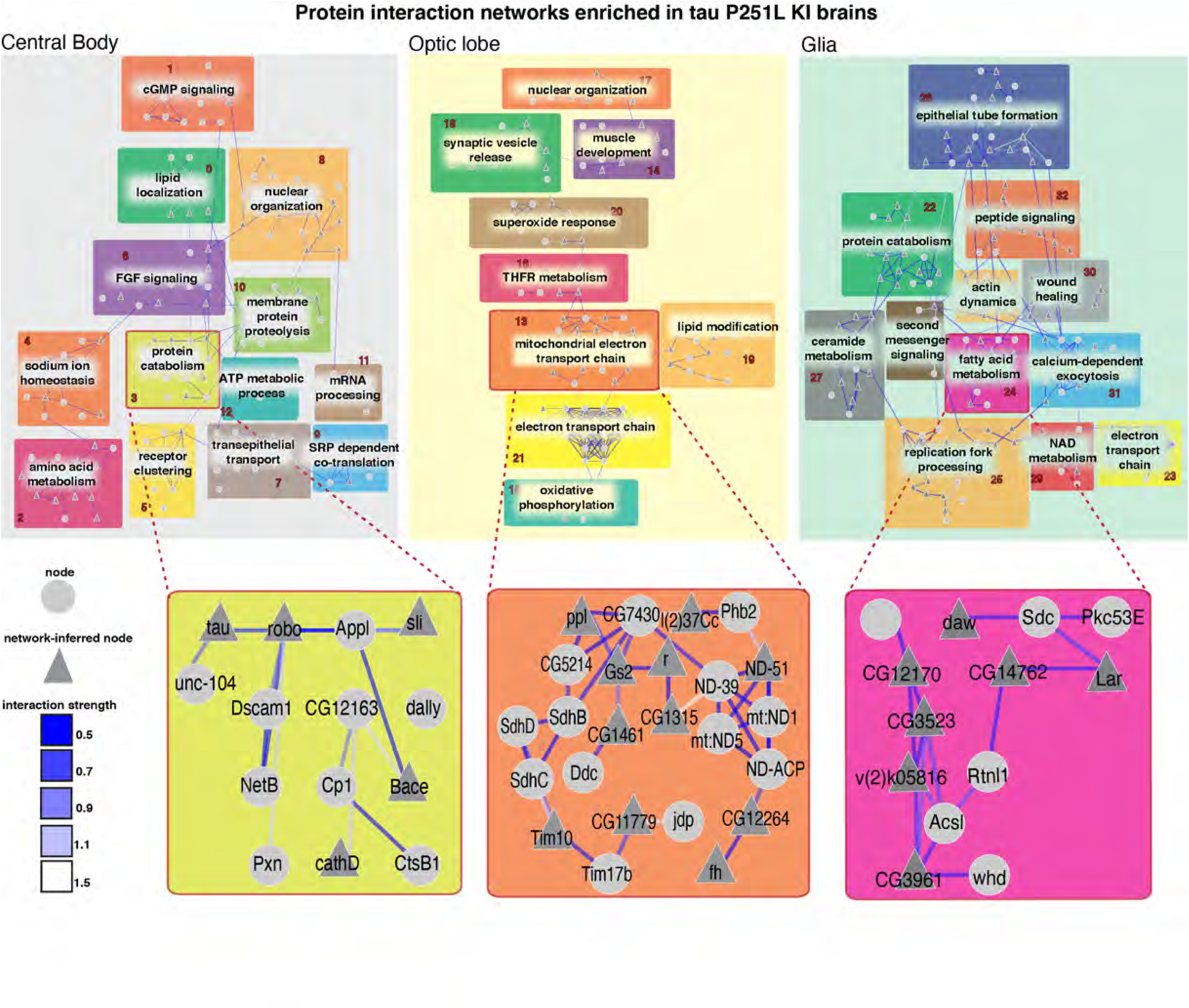
Protein interaction networks enriched in the central body, optic lobe and glia in tau P251L knock-in brains compared to controls. Protein interaction networks are largely distinct among central body neurons, optic lobe neurons and glia. Subnetworks including nodes enriched for protein catabolism (central body), electron transport chain (optic lobe) or fatty acid metabolism (glia) are highlighted. Interaction strength displayed in blue shows the stringency of the interaction: the lower the strength, the stronger the interaction.

Protein catabolism was a subnetwork in both central body and glial networks. Protein catabolism was connected to multiple other subnetworks in the central body network and interestingly contained multiple proteins previously implicated in Alzheimer’s disease, including Appl (fly ortholog of APP), beta-site APP-cleaving enzyme (Bace, a fly homolog of BACE1), three members of the cathepsin family (CtsB1, cathD, CtsF/CG12163), and tau itself identified as a computational network-inferred node. As expected from gene ontology analysis (Supplemental Fig. 7G), multiple metabolic subnetworks were identified in the glial network, consistent with the role of glia in providing metabolic support to neurons (Nedergaard and Verkhratsky 2012; Verkhratsky et al. 2012). A subnetwork enriched for nodes associated with fatty acid metabolism was identified in the glial network (Fig. 5), correlating with the important role of glia in lipid metabolism and signaling in both flies and mammalian systems (Goodman and Bellen 2022; Lee et al. 2021). Detailed protein interaction networks identified in the central body, optic lobe and glia are shown in Supplemental Fig. S8-S10.

### Cell-cell communication and pseudotime trajectory analyses highlight the role of glial cells in tau P251L knock-in brains

Altered gene expression (Supplemental Fig. 7) and protein interaction networks (Fig. 5) in glia driven by neuronal-predominant expression of P251L mutant tau suggests perturbed intercellular communication in P251L knock-in brains. We therefore calculated the interaction scores for 196 manually curated ligand-receptor pairs using the FlyPhoneDB quantification algorithm (Liu et al. 2022) in tau P251L knock-in brains and controls. We found significant alterations predicted in major cellular signaling pathways (Fig. 6; Supplemental Fig. S11). Altered signaling is indicated in circle plots in Fig. 6 (A,C,E,G) by nodes representing a unique cell types and edges representing a communication event. The thickness of an edge reflects the interaction strength of the communication event. Dot plots in Fig. 6 (B,D,F,H) display the calculated score of selected ligand-receptor pairs from one cell type to another with the shading of the dot indicating the interaction score and the size of the dot the P value. Many of predicated signaling changes support altered communication between glia and neurons. For instance, synaptic plasticity signaling, assessed by expression of the ligand spatzle and kekkon receptors, was mainly driven by perineurial glia in the control brain. However, perineurial glial cells in tau P251L knock-in animals had reduced expression of the ligand spatzle 5 while recipient cells downregulated kekkon receptors (Fig. 6B). Similarly, expression of the JAK-STAT ligand upd2 was significantly downregulated in perineurial glia in tau P251L knock-in brains compared to controls, while the receptor dome was reduced in expression in widespread target neuronal clusters (Fig. 6D). Interestingly, there was a predicted upregulation of JAK-STAT signaling from mushroom body output neurons to a restricted set of neuronal clusters in brains of flies expressing P251L mutant tau (Fig. 6C). In contrast, predicted hippo signaling from mushroom body output neurons to perineurial glial based on decreased levels of the ligand ds and receptor fat was decreased in tau P251L knock-in brains compared to controls (Fig. 6E).

**Figure 6:**
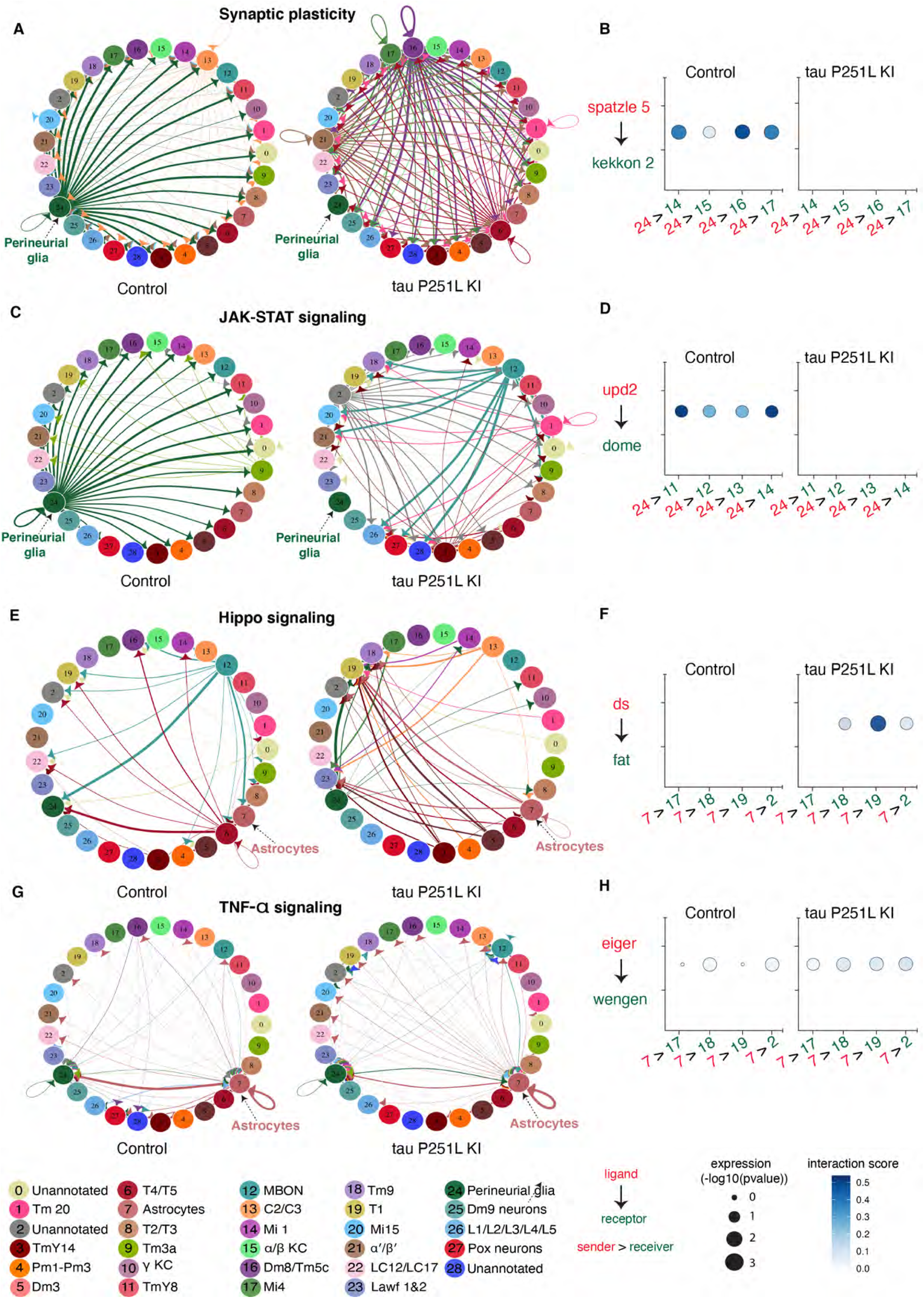
Cell-cell communication analysis predicts altered signaling in tau P251L knock-in brains compared to controls. Altered ligand and receptor expression predicts regulation of synaptic plasticity signaling mainly via perineurial glial cells in control brains (*A*). Signaling from perineurial glia is significantly reduced in tau P251L knock-in brains as predicted by levels of spaetzle ligand and kekkon receptor *(B)*. JAK-STAT signaling, as predicted by expression of the upd2 ligand and dome receptor, mediated by perineurial glia in control brains *(C)*, is substantially reduced in brains from tau P251L knock-in animals *(D)*. Hippo signaling, indicated by expression of ds ligand and fat receptor, is upregulated in astrocytes of flies expressing P251L mutant tau compared to controls *(E,F)*. Predicted TNF-α signaling from ligand eiger to receptor wengen is increased in astrocytes of tau P251L knock-in flies *(G,H)*. In panels (*B,D,F,H*) the interactions from and to the specified cell types are indicated on the x-axis, while the size of the circle indicates the P value and the intensity of the blue color illustrates the interaction score as defined in the figure label below the panels.

Astrocytic signaling also showed predicted changes in tau P251L knock-in brains compared to controls. JAK-STAT signaling perineurial glia to astrocytes was reduced in mutant tau expressing brains (Fig. 6C), while hippo signaling from astrocytes to multiple neuronal subtypes was increased in tau P251L knock-in brains (Fig. 6E,F). TNF-α signaling from astrocytes was also increased in flies expressing mutant tau, as suggested by increased levels of the ligand eiger and receptor wengen (Fig. 6G,H). Altered astrocyte integrin, hedgehog and insulin signaling was also suggested by changes in expression of ligand and cognate receptor pairs (Supplemental Fig. S11A,D,E).

Given altered gene expression (Fig. 4, Supplemental Fig. 7), protein interaction networks (Fig. 5) and predicted signaling (Fig. 6) in glia we next examined gene expression profiles in these non-neuronal cells in more detail (Fig. 7). Transposable elements were significantly upregulated in both types of glia (Fig. 7A,C; Supplemental Table S5), although one transposable element was highly downregulated in both glia subsets (RR48361). Gene ontology enrichment analysis highlighted different metabolic pathways in the two cell types. Amino acid and glutamate metabolism pathways were enriched in perineurial glia while L-cysteine, acyl-CoA and cAMP metabolic pathways were enriched in astrocytes.

**Figure 7:**
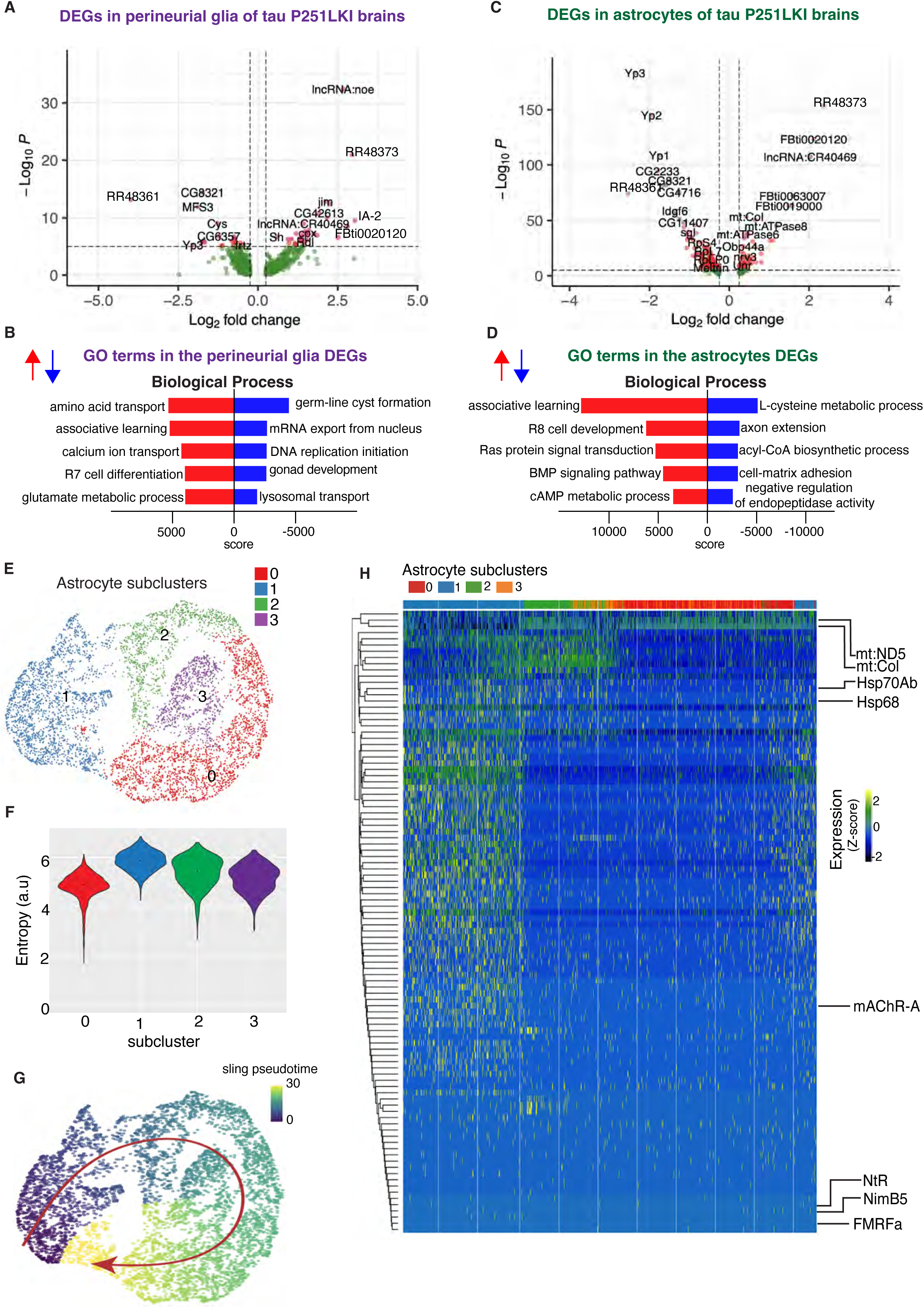
Gene expression and trajectory analysis in glia. Differentially regulated genes, both upreulated and downregulated, in perineurial glia of tau P251L knock-in brains compared to controls *(A)*. Gene ontology analysis shows biological processes associated with the up-regulated and down-regulated genes in perineurial glia from tau P251L knock-in brains compared to controls *(B)*. Differentially regulated genes, both upreulated and downregulated, in astrocytes of tau P251L knock-in brains compared to controls *(C)*. Gene ontology analysis shows biological process associated with upreulated and downregulated genes in astrocytes of tau P251L knock-in brains *(D)*. All dots on the volcano plots are significnat at FDR < 0.05 and log2FC > 0.25 for upregulated and < -0.25 for downregulated genes. Score represents the combined score c = log(p)*z (Chen et al. 2013). Astrocytes from both control and tau P251L knock-in brains were further subclustered into 4 groups. Entropy analysis to define the root for trajectory analysis revealed cluster 1 to have the highest entropy *(E,F)*. Slingshot trajectory analysis on astrocyte clusters identified a single lineage passing sequentially from clusters 1 to 2, 3, and 0 *(G)*. Differential gene expression between astrocyte subclusters adjacent in pseudotime were used to cluster genes along the pseudotime trajectory (*H*). Each row in the heat map represents a gene. The columns are astrocyte subclusters arranged arranged according to pseudotime from left to right. Examples of differentially regulated genes from enriched gene ontology biological processes are shown on the calculated trajectory (*H)*.

Since we observed significant alterations in glial signaling in tau P251L knock-in brains (Fig. 6; Supplemental Fig. S11) we investigated glial gene trajectories in our single-cell RNA sequencing, focusing on astrocytes because we obtained a large number (nearly 5800) of these cells (Supplemental Table S1). We first subclustered astrocytes into 4 groups (Fig. 7E). We then calculated the entropy of these clusters (Guo et al. 2017) and used cluster 1, which showed the highest entropy, as the root for trajectory analysis (Street et al. 2018). A single lineage starting from cluster 1 and progressing sequentially from cluster 2 through cluster 3 and finally to cluster 0 emerged (Fig. 7G). We then clustered differentially expressed genes along the calculated trajectory as presented in the heat map, in which pseudotime is represented in columns from left to right (Fig. 7H). Our pseudotemporal analysis suggests different stages of astrocytic response to tauopathy.

Gene ontology analysis across pseudotime revealed multiple genes involved in signaling pathways (*FMRFa, NimB5*), particularly in cholinergic signaling (nicotinic acetylcholine receptor subunit *NtR, mAChR-A, ChAT*) early in the glial trajectory. Cellular stress response emerged later in the trajectory with upregulation of heat shock proteins (*Hsp68*, *Hsp70Ab*), while altered mitochondrial gene expression (*mt:ND5, mt:Col*) characterized astrocytes late in the calculated trajectory. These findings suggest that altered astrocyte signaling (Fig. 6; Supplemental Fig. S11) may emerge early in tauopathy pathogenesis and drive subsequent cell-autonomous and non-cell autonomous stress responses and cytotoxicity. A complete list of all differentially expressed glial genes, genes associated with gene ontology biological processes, and trajectory-associated genes is provided in Supplemental Table S5.

### Gene regulatory networks in control and tau P251L knock-in Kenyon cells

Kenyon cells are a major defined neuronal component of the central body of the *Drosophila* brain (Fig. 4). Together with their output neurons (MBON), Kenyon cells play a central role in learning and memory in the *Drosophila* brain (Heisenberg 2003; Modi et al. 2020); memory loss is a key feature of human tauopathies (Grossman et al. 2023). Kenyon cells are cholinergic (Barnstedt et al. 2016), a neuronal type that is selectively vulnerable in previously described fly tauopathy models (Wittmann et al. 2001) and a pathway altered early in our trajectory analysis (Fig. 7). Our cell-cell communication analyses suggested altered signaling in Kenyon cells, or their output neurons, via multiple signaling pathways (Fig. 6; Supplemental Fig. 11). We therefore focused next on gene expression in Kenyon cells. We identified three Kenyon cells clusters, γ Kenyon cells, α/β Kenyon cells, and α’/β’ Kenyon cells (Fig. 8A). Transposable elements were upregulated in all Kenyon cell clusters in tau P251L knock-in brains (Supplemental Fig. S12A,C,E), as observed in other neuronal and glial clusters (Fig. 7, Supplemental Fig. 7). Analysis of biological pathways associated with common upregulated and downregulated genes in all three Kenyon cell clusters identified key biological processes previously linked to tauopathy pathogenesis (Götz et al. 2019; Frost et al. 2015), including control of DNA and RNA structure and metabolism (Fig. 8B), as well as many pathways without prior links to tauopathy. A complete list of differentially expressed genes and associated biological processes is given in Supplemental Table S6.

**Figure 8:**
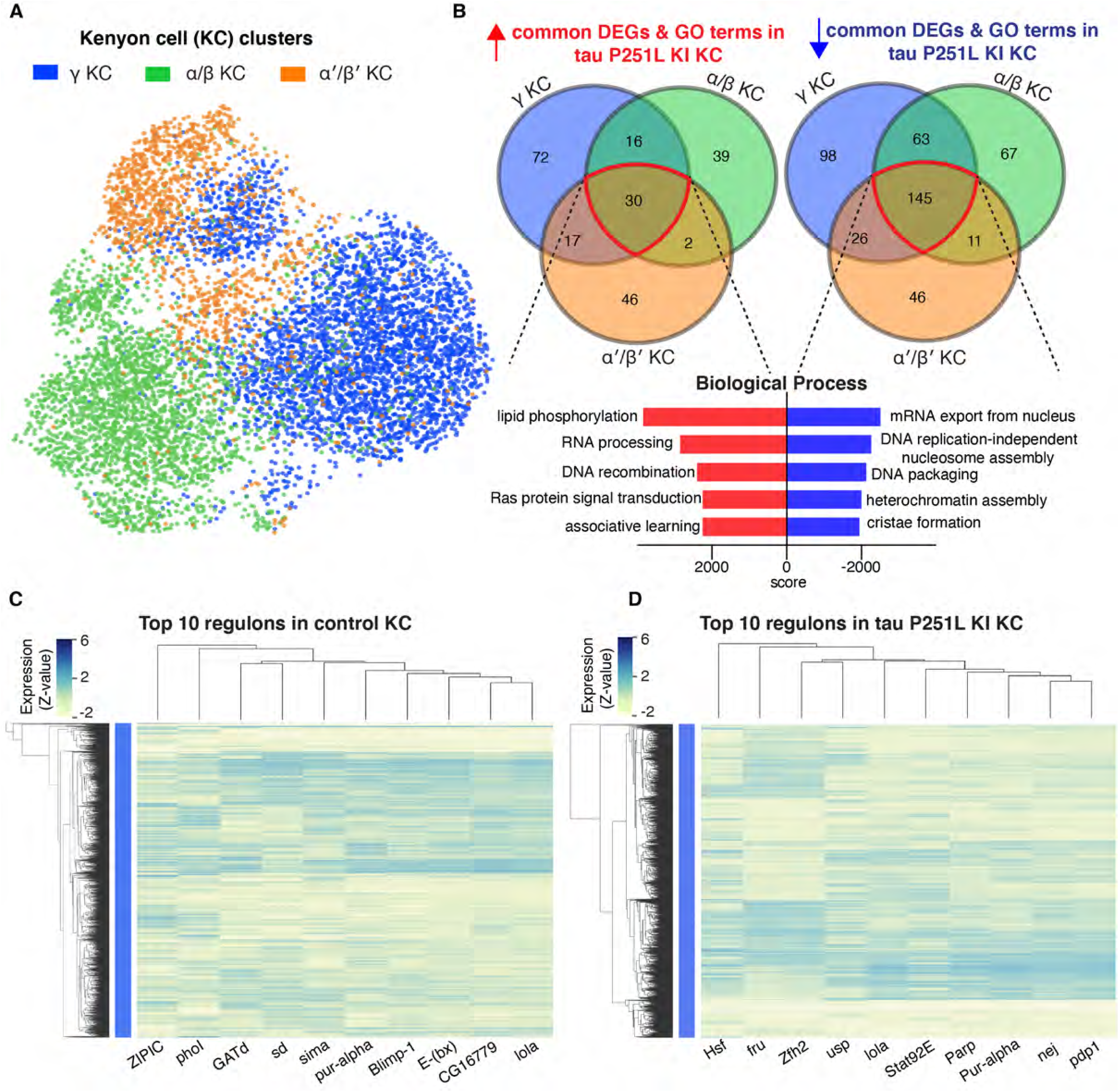
Gene expression and regulatory networks in Kenyon cells. Three Kenyon cell (KC) clusters, γ-KC, αβ-KC, and α’β’-KC, and biological process based on the common upreulated and downregulated genes in Kenyon cell clusters in tau P251L knock-in brains *(A,B).* Score represents the combined score c = log(p)*z (Chen et al. 2013). Control and tau P251L knock-in Kenyon cells were clustered separately using SCENIC gene regulatory network analysis to identify regulons. The top 10 regulons identified by SCENIC gene regulatory network analysis in control *(C)* and tau P251L knock-in *(C)* Kenyon cells *(C)* are presented in the heatmaps. Each row represents a Kenyon cell; each column is a regulon.

Given the multiple lines of evidence connecting tauopathy pathogenesis to Kenyon cell function we next determined the gene regulatory networks controlling disease-associated changes in gene expression in Kenyon cells. We implemented the SCENIC (Single-Cell rEgulatory Network Inference and Clustering, Aibar et al. 2017) workflow on gene expression data from control and tau P251L knock-in Kenyon cells. The top 10 regulons identified in control compared to tauopathy model Kenyon cells are show in columns in the heat maps in Fig. 8C (control Kenyon cells) and Fig. 8D (tau P251L knock-in Kenyon cells). Regulons were largely distinct in the two genotypes (Fig. 8C,D; Supplemental Table S7). The only shared transcription factor among the top 10 regulons was lola. Even for the shared lola regulon, the gene expression patterns per cell clustered and co-expressed with different transcription factors and are different among Kenyon cells of control vs. tau P251L knock-in animals. The distinct gene regulatory networks illustrated in the heatmap are concordant with altered gene expression (Fig. 8B) and cell-cell communication (Fig. 6) between control and tau P251L knock-in Kenyon cells. The increase in HSF, Stat92E and Parp expression (Supplemental Fig. 13) and regulons (Fig. 8D) in brains of tauopathy model flies are consistent with elevated cellular stress, DNA damage and cell death in aging neurons exposed to mutant tau P251L (Figs. 1,2).

## Discussion

Here we present a new model of tauopathy in the experimentally facile model organism *Drosophila* based on precise gene editing of the endogenous tau gene to introduce a mutation orthologous to human proline 301 to leucine (P301L), the most common tau mutation in frontotemporal dementia patients (Poorkaj et al. 2001). We observe age-dependent neurodegeneration in our knock-in animals (Fig. 1C,D). Homozygous knock-in flies display early and greater total levels of degeneration compared to heterozygous animals. These findings are compatible with a toxic gain of function mechanism, as generally posited in familial frontotemporal tauopathies (Goedert et al. 2012; Frost et al. 2015; Götz et al. 2019; Bardai et al. 2018b). However, given the important role of microtubules in neurodevelopment, a loss of function component contribution cannot be excluded, even given the lack of clear neurodegeneration in tau knockout mice (Harada et al. 1994; Dawson et al. 2001; Morris et al. 2013) and flies (Burnouf et al. 2016). As expected given that levels of mutant tau are controlled by the endogenous tau promotor in our model compared with the strong exogenous promotor systems employed in prior transgenic models, neurodegeneration in knock-in animals is observed at older ages and is milder (Wittmann et al. 2001; Bardai et al. 2018b; Law et al. 2022). However, we do observe key biochemical and cellular pathologies previously described in transgenic *Drosophila* tauopathy models, including metabolic dysfunction (Fig. 2A,B), elevated levels of DNA damage (Fig. 2C-G), and abnormal cell cycle activation (Fig. 1E) (Khurana et al. 2012; DuBoff et al. 2012; Bardai et al. 2018a; Khurana et al. 2006). Importantly, these pathways are also perturbed in mouse tauopathy models and tauopathy patients (Khurana et al. 2012; Götz et al. 2019; Frost et al. 2015; Andorfer et al. 2005; DuBoff et al. 2013; Welch and Tsai 2022; Herrup and Arendt 2002).

The similarities of our knock-in model to human tauopathies and prior overexpression tauopathy models, recapitulated in a more faithful genetic knock-in context, motivated us to perform a comprehensive transcriptional analysis in our tau P251L knock-in brains using single-cell RNA sequencing. We recovered a large number (130,489) of high-quality cells, which allowed us to identify the majority of previously annotated neuronal and glial groups from prior single cell sequencing analyses in the adult fly brain (Li et al. 2022; Davie et al. 2018). Comparing gene expression profiles between control and tau P251L knock-in animals revealed pervasive dysregulation of genes in neuronal (Figs. 4,8) and glial (Fig. 7) subtypes and throughout different anatomic regions (Fig. 4, and Supplemental Fig. 6). These findings are consistent with prior single cell sequencing studies in flies overexpressing mutant human tau (Praschberger et al. 2023; Wu et al. 2023). We observed regulation of both common and distinct biological pathways when comparing differentially expressed genes across cell subtypes. Transposable elements were notably upregulated in the complete gene expression set, as well as in specific anatomic regions and neuronal subtypes. These findings correlate with a previously described functional role for transposable element mobilization in *Drosophila* models of tauopathy, and in tauopathy patients (Sun et al. 2018; Guo et al. 2018). Mitochondrial function has been strongly linked to neurotoxicity in tauopathies (Frost et al. 2015; Götz et al. 2019; DuBoff et al. 2013) and is a feature of our current model (Fig. 2). We accordingly observed altered expression of mitochondrial genes and biological processes in the complete expression data set (Fig. 4), as well as in separate analyses of the central body, optic lobe (Supplemental Fig. 7), and Kenyon cells (Fig. 8, Supplementary Fig. S12). More importantly, we observed significant alterations in multiple metabolic, cellular communication and biological pathways not previously implicated in tauopathy pathogenesis (Figs. 4,5,6), which can now be assessed in tauopathy models and patients for mechanistic relevance and ultimately therapeutic targeting.

Cell type selectivity is a fundamental, and poorly understood, feature of human neurodegenerative diseases, including tauopathies. Our protein interaction networks highlighted regionally specific biology with predominantly distinct nodes appearing in the central body compared to the optic lobe (Fig. 5). Comparative analysis of genes differentially expressed in central body compared to the optic lobe are consistent with substantial regional differences in the response to mutant tau expression with substantially greater numbers of unique compared to common genes upregulated in the central body vs. the optic lobe (Supplemental Fig. 7E). Even within subgroups of Kenyon cells there are equivalent numbers or more uniquely up- or down-regulated genes compared to commonly regulated genes (Fig. 8B). Our dataset thus highlights a substantial set of genes that may contribute to selective neuronal susceptibility in neurodegeneration, including many differentially regulated genes and processes not previously linked to tau pathobiology.

Although tau is a predominantly neuronal protein (Götz et al. 2019; Goedert 2004; Heidary and Fortini 2001), we observed significant alteration of glial gene expression in tau P251L knock-in brains compared to controls (Figs. 4,7), suggestive of non-cell autonomous control of glia cell function by neuronally expressed tau. Gene ontology (Fig. 7A,B) and protein interaction network (Fig. 5) analyses highlighted a number of metabolic processes altered in glia by expression of toxic tau in neurons, including glutamate, lipid and amino acid metabolism (Figs. 5,7). Glial uptake and detoxification of neurotransmitters and their metabolites, as well as toxic lipid species, maintains neuronal function and viability. Lipid metabolism is further central to energy production by glial cells, which support highly energy consuming neurons with active synaptic transmission (Smolič et al. 2021; Jiwaji and Hardingham 2023). In addition to glial processes previously implicating in controlling neuronal health, our transcriptional analysis revealed new metabolic and signaling pathways in glia regulated by expression of mutant tau (Fig. 7A-C), which can now be explored as non-cell autonomous mechanisms regulating neuronal function and viability in tauopathy.

An effect of mutant tau expression in neurons on glial gene expression implies signaling, and possibly perturbed signaling, between the two cell types. Examination of expression of 196 ligand-receptor pairs (Liu et al. 2022) indeed supported broad alterations in glial-neuronal communication in tau P251L knock-in flies (Fig. 6, Supplemental Fig. S11), with mutant tau expression perturbing synaptic plasticity, JAK-STAT, hippo, TNF-α, integrin and EGFR signaling between perineurial cells, astrocytes and multiple neuronal subtypes. Although prior studies have implicated glial signaling, for example the JAK-STAT pathway (Colodner and Feany 2010), in non-cell autonomous control of neurotoxicity, the pervasive nature of the altered signaling suggested by our single-cell transcriptional analyses is unexpected and provides multiple targets for functional testing. Our findings further suggest that a systematic and broad perturbation of intercellular signaling is present in tauopathy, which may require manipulation of multiple pathways to correct and systems-level analysis to monitor.

Trajectory analysis has been widely used to order temporal events along developmental pathways, but has less often been applied to neurodegenerative disease progression (Karademir et al. 2022; Wang et al. 2022; Fitz et al. 2021; Dai et al. 2023). Given the evidence for altered glial-neuronal communication in our tau knock-in model we assessed possible trajectories in the four distinct subgroups of astrocytic glial cells that we defined. Using the astrocyte cluster with the highest entropy as the root (Guo et al. 2017) we identified a single astrocyte trajectory (Fig. 7G). Differential gene expression and gene ontology analyses across the trajectory revealed altered expression of neurotransmitter and cell signaling genes first, followed by altered cell stress responses, and finally mitochondrial changes (Fig. 7H, Supplemental Table S5). A number of genes involved in cholinergic signaling were changed early in the glial trajectory. We have previously demonstrated that cholinergic terminals are preferentially vulnerable and degenerate early in a tauopathy model based on transgenic human tau expression in flies (Wittmann et al. 2001). Our trajectory analysis may thus help identify early events in glial-mediated neurodegeneration, including pathways not previously associated with tauopathy (Supplemental Table S5). Glial pathways contributing to neurodegeneration are increasingly recognized as attractive and understudied avenues for therapeutic intervention (Jiwaji and Hardingham 2023). Identifying and intervening in early glial-neuronal signaling events may prevent later, and possibly irreversible, neuronal damage.

Reversing pathological neuronal cell-autonomous programs may provide an alternative or additional method of preventing neuronal dysfunction and death in tauopathies. We focused on Kenyon cells as a group of neurons involved in the behaviorally relevant process of memory and comprised of cholinergic neurons, a vulnerable cell type in *Drosophila* (Wittmann et al. 2001) and human (Ishida et al. 2015; Whitehouse et al. 1981) tauopathies to define transcriptional programs driving neurodegeneration in response to mutant tau expression. As expected by the multiple neuropathological and cell biological abnormalities present in our knock-in model flies (Figs. 1,2), we observed substantially distinct regulons in tau P251L knock-in Kenyon cells compared to controls (Fig. 8C,D). We identified regulons involved in stress responses (Hsf, Stat92E), including the DNA damage response (Parp), as would be expected from the presence of elevated DNA damage in Kenyon cells in our knock-in flies (Fig. 2E-G). We recovered nej, the fly ortholog of vertebrate CREB-binding protein (CBP) as a top regulon induced in knock-in flies. Increasing levels of nej/CBP is beneficial in fly (Cutler et al. 2015) and vertebrate (Caccamo et al. 2010) models relevant to Alzheimer’s disease, suggesting that upregulation of nej may represent a protective response in Kenyon cells. We also identified multiple regulons not previously associated with neurodegenerative tauopathies (Fig. 8C,D). Therapeutic manipulation of these programs or key transcriptionally regulated mediators will be attractive candidates for evaluation in patient tissue, patient derived cellular models and vertebrate models of tauopathy.

The mechanisms transducing the effects of mutant tau on gene expression are likely multiple and as yet only partially characterized. We have previously defined a cascade in which cytosolic tau binds and stabilizes F-actin (Fulga et al. 2007), leading to signal transduction through the LINC complex, nuclear lamin disruption (Frost et al. 2016) and consequent chromatin relaxation (Frost et al. 2014) promoting aberrant transposable element activation and neurodegeneration (Sun et al. 2018). Other cytosolic targets of tau may promote transcriptional regulation through parallel mechanisms. For example, tau-mediated actin hyperstabilization promotes mitochondrial dysfunction and excess production of oxidative free radicals by interfering with mitochondrial dynamics (DuBoff et al. 2012). Oxidative stress may directly contribute to elevated DNA damage in tauopathy (Bardai et al. 2018b; DuBoff et al. 2013; Götz et al. 2019; Frost et al. 2016). However, although tau is best known as a cytosolic protein involved in regulation of the cytoskeleton, a number of studies have demonstrated that tau can also be detected in the nucleus (Loomis et al. 1990; Thurston et al. 1996; Cross et al. 2000), where the protein binds DNA (Wei et al. 2008; Hua et al. 2003; Sjöberg et al. 2006; Bukar Maina et al. 2016). Thus, tau may play a direct role in instructing the nuclear transcriptional programs we have defined (Fig. 8C,D).

In summary, here we develop a genetically precise model of frontotemporal dementia caused by the most common tau mutation found in patients and present a comprehensive picture of gene expression changes and derived protein interaction, cell signaling and transcriptional networks. We recapitulate neurodegeneration, metabolic dysfunction and DNA damage, common features of human tauopathies (Goedert 2004; Götz et al. 2019; Welch and Tsai 2022) and confirm that cellular pathways perturbed in overexpression tauopathy models are also dysregulated in the more faithful genetic knock-in context. More importantly, our work suggests previously unsuspected, pervasive alterations in glial-neuronal signaling in tauopathy pathogenesis, implicates many new genes and pathways and provides a genetic model system in which to test the new hypotheses our data suggests.

## Methods

### Genetics and CRISPR-Cas9 editing

The *Drosophila tau* gene is located on the 3rd chromosome. The guide RNAs targeting the *tau* gene to mutate proline 251 to leucine were identified using Harvard Medical School’s DRSC/TRiP “find CRISPRs” tool. The gRNA ’5 CCGGGAGGCGGGGACAAGAAGAT 3’ was cloned into pCDF3.1 plasmid and injected into the embryos of the TH_attP40 nos-Cas9 strain along with a single-stranded oligo nucleotide donor. The single-stranded oligo nucleotide donor was 150 bp in length and contained a C to T transition that resulted in alteration of the codon CCG (proline) to CTG (leucine). Embryos were injected (BestGene Inc.) and founder flies obtained. Founder flies were then balanced to obtain homozygous knock-in animals. The mutation was confirmed by PCR. The genotype of knock-in animals in most experiments (Figures 1,2C-E,4-8) was *elav-GAL4/+; tau-P251L knock-in* (homozygous or heterozygous for *tau-P251L knock-in* as specified in figures and legends). In these experiments control animals were *elav-GAL4/+.* In Fig. 2A,B the genotype of knock-in flies was *w^1118^; tau-P251L knock-in* / *tau-P251L knock-in* (homozygous) or *w^1118^; tau-P251L knock-in* / *+* (heterozygous) as specified in the figure. In Fig. 2A,B the genotype of control flies was *w^1118^*. The *elav-GAL4* line was obtained from the Bloomington *Drosophila* Stock Center. Patrik Verstreken kindly provided tau knockout flies. All crosses and aging were performed at 25°C.

### Assessment of neurodegeneration and metabolism

For sectioning, adult flies were fixed in formalin at 1, 10 and 30 days of age and embedded in paraffin. Vacuoles, PCNA and pH2Av levels were examined using previously described methodology (Fulga et al. 2007; Frost et al. 2014) with additional details provided in the Supplemental Methods. Primary antibodies used include pH2Av (Rockland, 600-401-914, 1:100), elav (DSHB, 9F8A9, 1:5), GAPDH (Thermo Fisher, MA5-15738, 1:1000) and PCNA (DAKO, MO879, 1:500). A polyclonal antibody to *Drosophila* tau was prepared in rabbits immunized with full length recombinant tau protein (Thermo Fisher) and was used at 1:5,000,000 for western blotting. For all histological analyses, at least 6 brains were analyzed per genotype and time point. The comet assay and assessment of bioenergetics were performed as previously described (Frost et al. 2014; Sarkar et al. 2020) with additional details provided in the Supplemental Methods. The sample size (n), mean and SEM are given in the figure legends. All statistical analyses were performed using GraphPad Prism 5.0. For comparisons across more than 2 groups, one-way ANOVA with Tukey post-hoc analysis was used. For comparison of 2 groups Student’s t-tests were performed.

### Single-cell RNA sequencing (scRNA-seq) and downstream analyses

A standard sample preparation (Li et al. 2017; Davie et al. 2018), raw data processing (Satija et al. 2015) and downstream analyses such as cell cluster annotation (Hu et al. 2021) gene ontology analysis (Kuleshov et al. 2016), protein-protein interaction network analysis (Tuncbag et al. 2016), cell-cell communication analysis (Liu et al. 2022), trajectory analysis (Street et al. 2018) and gene regulatory network analysis(Van de Sande et al. 2020) were performed as previously described. Detailed methods are presented in the Supplemental Methods.

## Data Access

All raw and processed sequencing data generated in this study have been submitted to the NCBI Gene Expression Omnibus (GEO; https://www.ncbi.nlm.nih.gov/geo/) under accession number GSE223345. R code that was used to perform Seurat-based integration, trajectory, and cell-cell interaction and PPI network analyses are available at GitHub (https://github.com/bwh-bioinformatics-hub/Single-cell-RNA-seq-of-the-CRISPR-engineered-endogenous-tauopathy-model) and in the Supplemental Code file.

## Competing Interest Statement

The authors have no competing interests.

## Acknowledgments

We thank Tingting Zhao for help with bioinformatics analyses and Yi Zhong for excellent technical assistance. Fly stocks obtained from the Bloomington *Drosophila* Stock Center (NIH P40OD018537) were used in this study. We thank Dr. Patrik Verstreken for providing the *Drosophila* tau knockout line. Monoclonal antibodies were obtained from the Developmental Studies Hybridoma Bank developed under the auspices of the NICHD and maintained by the University of Iowa, Department of Biology, Iowa City, IA 52242. This research was funded by NIH R01AG057331 and AG076214 and Aligning Science Across Parkinson’s [Grant number ASAP-000301] through the Michael J. Fox Foundation for Parkinson’s Research (MJFF). For the purpose of open access, the author has applied a CC BY public copyright license to all Author Accepted Manuscripts arising from this submission.

## Notes

### Competing Interest Statement

The authors have declared no competing interest.

