## Supplemental Figures for "Transcriptional programs mediating neuronal toxicity and altered glial-neuronal signaling in a *Drosophila* knock-in tauopathy model"

```
H. Sapiens          MAEPRQEFEVMEDHAGTYGLGDRKDQGG-YTMHQDQEGD---TDAGLKESPLQTPTEDGSEE 58
M. Musculus         MADPRQEFDTMEDHAGDY------------TLLQDQEGD---MDHGLKESPPQPPADDGAEE 47
B. Tarus            MAEPRQEFDVMEDHAQ----------GD-YT-LQDQEGD---MDPGLKESPLQTPADDGSEE 47
C. Elegans(PTL1)    MSTPQSEPGSEPGSP-----------GENENSMETQYSQGSFPEVAVEPEPEPEP------E 45
D. Melanogaster     MAD-VLEKSSLLDAVPP--------LGDPHPPLPHQQLQ---QEAAAA-------------- 36

H. Sapiens          PGSETSDAKSTPTAEDVTAPLVD---EGAPGKQAAAQP-HTEIPEGTTAEEAGIG-DTPSLE 115
M. Musculus         PGSETSDAKSTPTAEDVTAPLVD---ERAPDKQAAAQP-HTEIPEGITAEEAGIG-DTPNQE 104
B. Tarus            PGSETSDAKSTPTAEDATAPLVD---EGAPGEQAAAQA-PAEIPEGTAAEEAGIG-DTSNLE 104
C. Elegans(PTL1)    PEPE-------PEPE----P------EPEP--------EPEPKPK---PEVEPVPQPEPEPE 79
D. Melanogaster     ----------------------------------------------AAANAA-PPAPPQQ 49

H. Sapiens          DEAAGHVTQARMVSKSKDGTGSDDKKAKGADGKT--KIATPRGAAPPG--QKGQANATRIPA 173
M. Musculus         DQAAGHVTQARVASKD--RTGNDEKKAKGADGKTGAKIATPRGAASPA--QKGTSNATRIPA 162
B. Tarus            DQAAGHVTQARMVSKGKDGTGPDDKKTKGADGKPGTKIATPRGAAPPG--QKGQANATRIPA 164
C. Elegans(PTL1)    YQPEPEPEPEVEPEPEPEPVPEPEPEP-EPEPEPVVEKEEEVVVESPPREQEPEKSGKSKPS 140
D. Melanogaster     QQPPPHQLQQQPQQQQLQQKPANARANQD-QKEG---DNDSGVDEST--QEKDRNGPNSPS 105

H. Sapiens          KTPPAPKT----------PP--SS----GEPPKSGDRSGYSSP------------------- 200
M. Musculus         KTTPSPKT----------PP--GS----GEPPKSGERSGYSSP------------------- 189
B. Tarus            KTTPTPKTSPATMQVQKKPPPAGAKSERGESGKSGDRSGYSSP------------------- 207
C. Elegans(PTL1)    SPIPDA------------PTMEDIAPRELESLNFSETSGTSDQQADRIMQNNENERVEEKKQ 190
D. Melanogaster     SPV-KT------------PT–ST-----SSKP---DKSGTSRP------------------- 126

H. Sapiens          --GSPGTPGSRSRTPSLPTPPT---REPKK-VAVVRTP----PK-SPSSAKSRLQTA---PV 248
M. Musculus         --GSPGTPGSRSRTPSLPTPPT---REPKK-VAVVRTP----PK-SPSASKSRLQTA---PV 237
B. Tarus            --GSPGTPGSRSRTPSLPTPPT---REPKK-VAVVRTP----PK-SPSAAKSRLQAA---PG 255
C. Elegans(PTL1)    MSPTPSQPQHKTPQRSGIRPPTAILRQPKPIPASLPRPATAT-PSSQRAISTPRQTASTAPS 251
D. Melanogaster     --PS-ATPSNKS------APKS---RSASKNRLLLKTPE---PEPVKKVPMNKVQVG---HA 170

                                                     MTBD repeat
H. Sapiens          PMPD--------------------------------LKNVKSKIGSTENLKHQPGGG------ 273
M. Musculus         PMPD--------------------------------LKNVRSKIGSTENLKHQPGGG------ 262
B. Tarus            PMPD--------------------------------LKNVKSKIGSTENLKHQPGGG------ 280
C. Elegans(PTL1)    PRPISKMSRERSDVQKSTSTRSIDNVGRMTPKVNAKFVNVKSKVGSVTN--HKAGGGNVEIF 311
D. Melanogaster     PSPN--------------------------------LKAVRSKIGSLDNATYKPGGGHVKIE 200

                                                      MTBD repeat
H. Sapiens          --------------------------KVQIINKKLDLSNVQSKCGSKDNIKHVPGGGSVQIV 309
M. Musculus         --------------------------KVQIINKKLDLSNVQSKCGSKDNIKHVPGGGSVQIV 298
B. Tarus            --------------------------KVQIINKKLDLSNVQSKCGSKDNIKHVPGGGSVQIV 316
C. Elegans(PTL1)    SEKRLY-NAQSKVGSLKNATHVAGGGNVQIENRKLDFSAASPKVGSKTN--YQPAKSDVKIV 370
D. Melanogaster     SKKIDIKAAPRIEA--KNDKYMPKGGEKKIVTTKL-QWNAKSKIGSLENAAHKPGGGDKKIE 259

                       MTBD repeat                          MTBD repeat
H. Sapiens          YKPVDLSKVT-SKCGSLGNIHHKPGGGQVEVKSEKLDFKDRVQSKIGSLDNITHVPGGGNKK 370
M. Musculus         YKPVDLSKVT-SKCGSLGNIHHKPGGGQVEVKSEKLDFKDRVQSKIGSLDNITHVPGGGNKK 359
B. Tarus            YKPVDLSKVT-SKCGSLGNIHHKPGGGQVEVKSEKLDFKDRVQSKIGSLDNITHVPGGGNKK 316
C. Elegans(PTL1)    SEKL-TWQAK-SKVGSMDNAAHKPAGGNVQILSQKL-NW-KAESKVGSKDNMNHRPGGGNVQ 428
D. Melanogaster     TLKMDFKDKAKPKVGSTANVKHQPGGGDIKIQTQKLEIKA--QSKVGSLDNVKHKPGGGEKK 319

H. Sapiens          IETHKLTFRENAKAKTDHGAEIVYKSPVVSGDTSPRHLSNVSSTGSIDMVDSPQLATLADEV 432
M. Musculus         IETHKLTFRENAKAKTDHGAEIVYKSPVVSGDTSPRHLSNVSSTGSIDMVDSPQLATLADEV 421
B. Tarus            IETHKLTFRENAKAKTDHGAEIVYKSPVVSGDTSPRHLSNVSSTGSIDMVDSPQLATLADEV 439
C. Elegans(PTL1)    IFDEKIRYVSTDSSRNHSTLDISSL------------------------------------- 453
D. Melanogaster     IFDDKDYLKNVEHSVALTTPPTQSPLPSMTASGADENLNQQS-------------------- 361

H. Sapiens          SASLAKQGL    441
M. Musculus         SASLAKQGL    430
B. Tarus            SASLAKQGL    448
C. Elegans(PTL1)    ---------    453
D. Melanogaster     ---------    361
```

**Supplemental Fig. S1: Multiple sequence alignment of tau across species.** Alignment of human tau (4 repeat) and C. elegans PTL1 (tau homolog) to tau from different species. Conserved residues are highlighted in bold, and human proline 301 and orthologous prolines in other species are shown in bold red. The alignment highlights conservation of microtubule binding domains (MTBD).

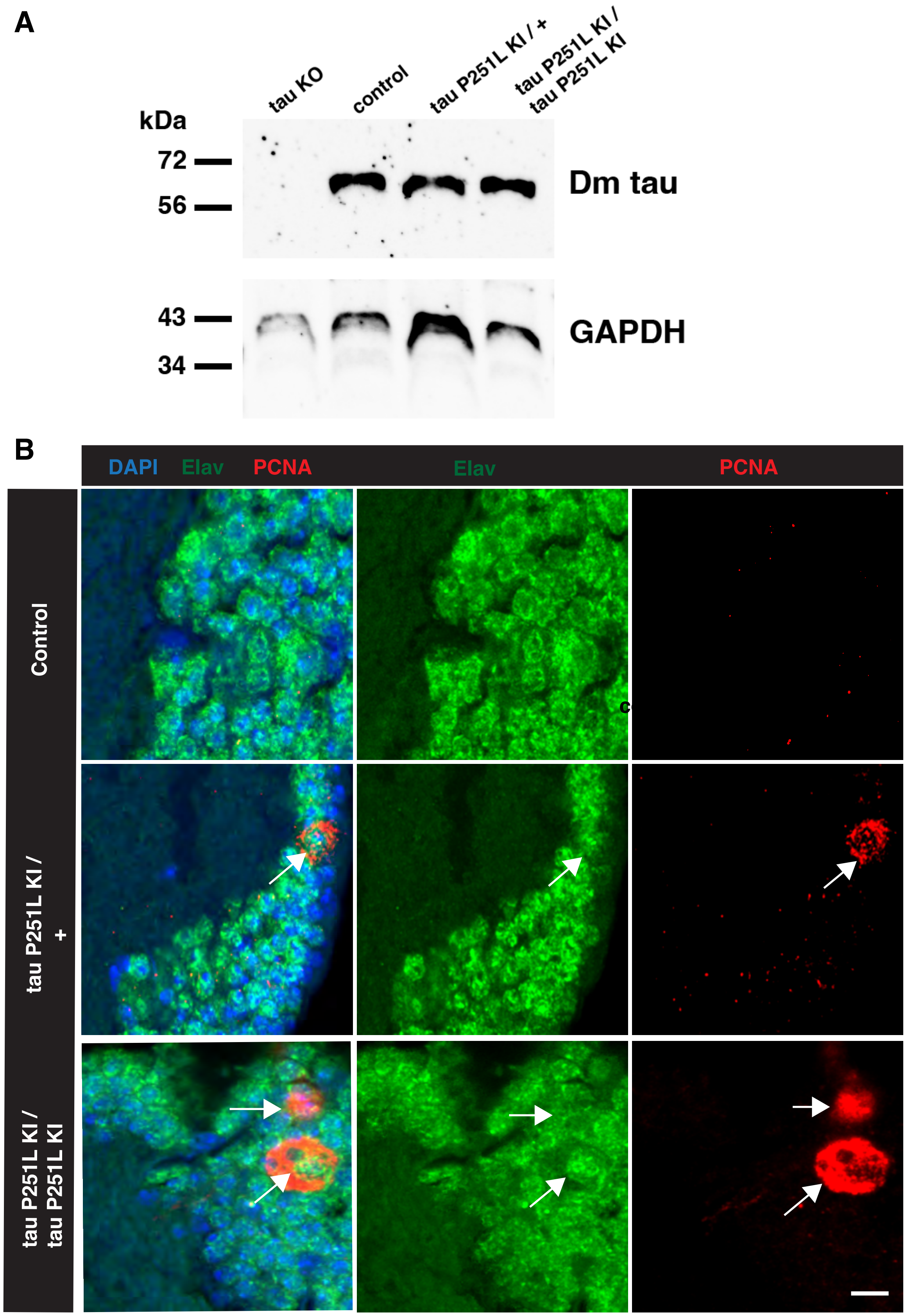


Supplemental Fig. S2: *Drosophila* tau levels and PCNA staining in tau P251L KI brains compared to controls. A) Immunoblotting analysis using an antibody to *Drosophila* tau shows equivalent levels of wild type and P251L tau. The blot is reprobed with an antibody to GAPDH to illustrate equivalent protein loading. B) Representative images of proliferating cell nuclear antigen staining in control and tau P251L knock-in Kenyon cells (arrows, identified with the neuronal marker elav). Control is *elav-GAL4/+*. Flies are 10 days old in (A) and 30 days old in (B).

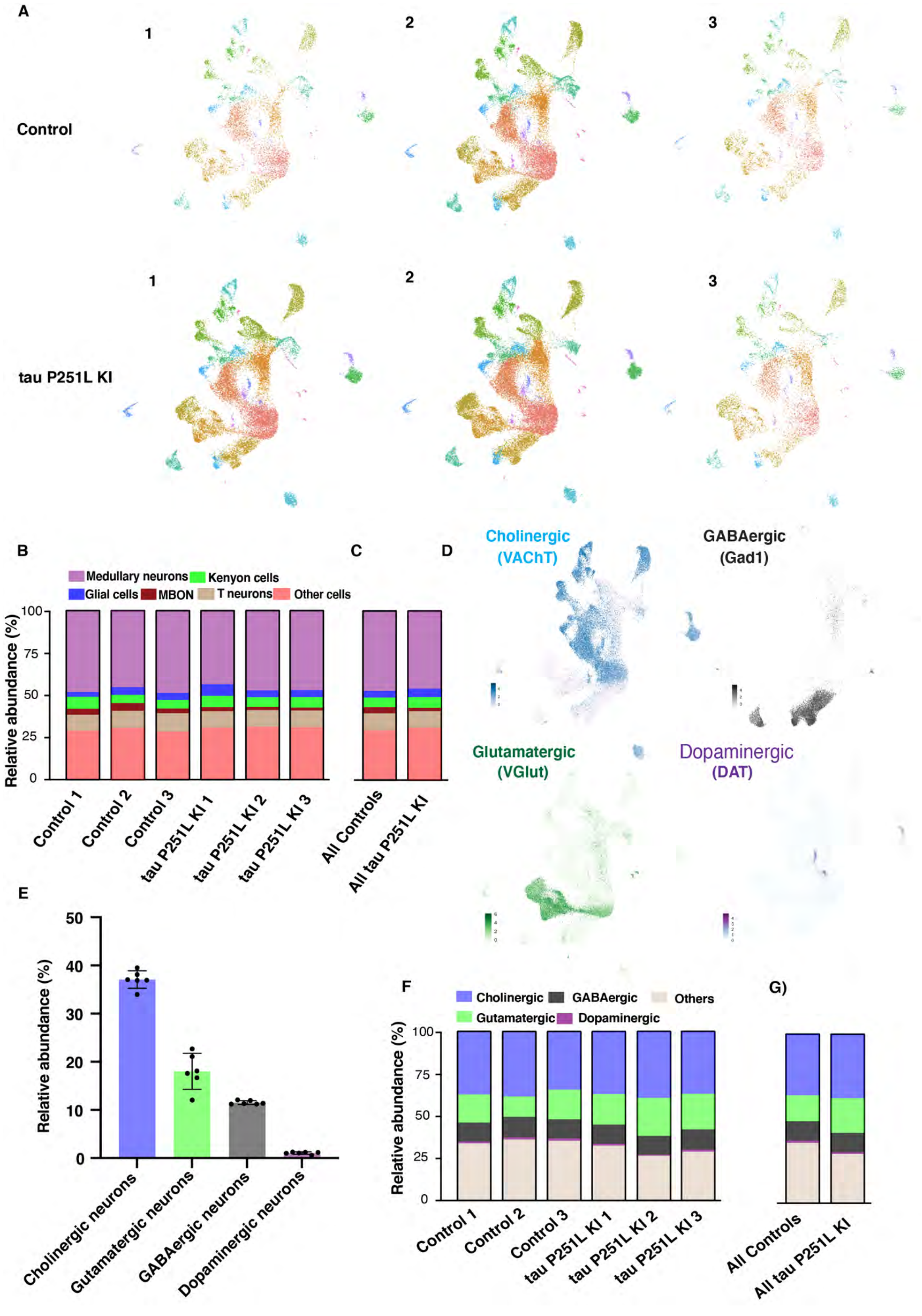
A
1
2
3
Control
1
2
3
tau P251L KI
B
C
D
Medullary neurons
Kenyon cells
Glial cells
MBON
T neurons
Other cells
Relative abundance (%)
100
75
50
25
0
Control 1
Control 2
Control 3
tau P251L KI 1
tau P251L KI 2
tau P251L KI 3
All Controls
All tau P251L KI
Cholinergic
(VAChT)
GABAergic
(Gad1)
Glutamatergic
(VGlut)
Dopaminergic
(DAT)
E
50
40
30
20
10
0
Relative abundance (%)
Cholinergic neurons
Gutamatergic neurons
GABAergic neurons
Dopaminergic neurons
F
Cholinergic
GABAergic
Others
Gutamatergic
Dopaminergic
G)
Relative abundance (%)
100
75
50
25
0
Control 1
Control 2
Control 3
tau P251L KI 1
tau P251L KI 2
tau P251L KI 3
All Controls
All tau P251L KI

**Supplemental Fig. S3: Relative abundance of the cells and neuronal identities within the final integrated dataset.** Individual 10x runs were integrated and contained cells from all clusters: 3 of control and 3 of tau P251L KI and relative abundance (%) of the medullary neurons (14 clusters), Kenyon cells (3 clusters), glial cell (2 clusters), MBON cluster, and T-neurons (3 clusters) in the individual sc-RNA seq runs *(A&B)*. Relative abundance (%) of the 3 integrated control and tau P251L KI datasets *(C)***.** Relative expression (>2 fold) of the key neuronal identity markers genes, such as VAChT for cholinergic neurons, Gad1 for GABAergic neurons, VGlut for glutamatergic neurons, and DAT for dopaminergic neurons, within the integrated dataset *(D)*. Relative abundance (%) of the neuronal identities within the integrated dataset *(E)*. Relative abundance (%) of the neuronal identities in the individual sc-RNA seq runs *(F)*. Relative abundance (%) of neuronal identities in the 3 integrated control and tau P251L KI datasets *(G)***.**

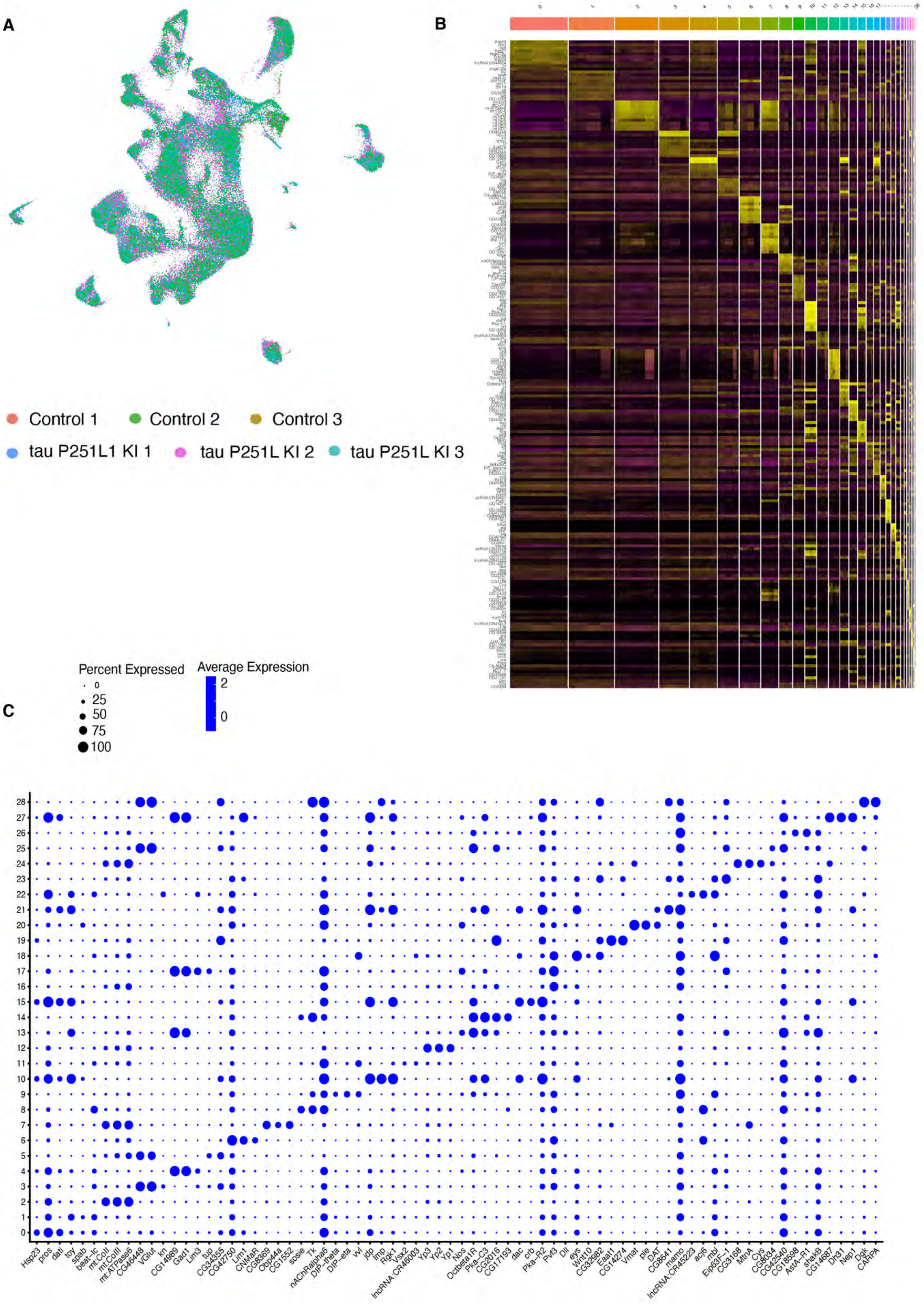
A
B
C
Control 1
Control 2
Control 3
tau P251L1 KI 1
tau P251L KI 2
tau P251L KI 3
Percent Expressed
0
25
50
75
100
Average Expression
2
0
Hsp23 pros dati toy spab beat-Ic mt:CoI mt:CoIII mt:ATPase6 CG46448 VGlut kn CG14989 Gad1 Lim3 tup CG34355 CG42750 Lim1 CNMaR CG8369 Obp44a CG1552 sosie Tk nAChRalpha6 DIP-theta DIP-eta vvl jdp Imp Rgk1 Vsx2 lncRNA:CR46003 Yp3 Yp2 Yp1 Nos Octbeta1R Pka-C3 CG2016 CG17193 dac crb Pka-R2 Pvf3 Dll eys Wnt10 CG32982 Eaat1 CG14274 Vmat ple DAT CG8641 mamo lncRNA:CR45223 acj6 mbl Eip63F-1 CG3168 MtnA Cys CG8034 CG42540 CG18598 AstA-R1 shakB CG14687 Dh31 Nep1 Dgk CARPA

**Supplemental Fig. S4: Expression profile of the top genes within the integrated dataset. (A)** The proportion of each sample, 3 of control and 3 of tau P251L KI, within the final integrated dataset obtained after single-cell RNA seq bioinformatics pipeline *(A)*. Heatmap of the top 10 highly expressed genes emerged after dimensionality reduction in all the clusters within the integrated dataset *(B)*. A dot plot showing the percentage expression of the top 3 genes within each single cell cluster identified in the integrated dataset. These 3 top markers and other top 7 markers (top 10 markers) were used to annotate the single cell cluster identities *(C)*.

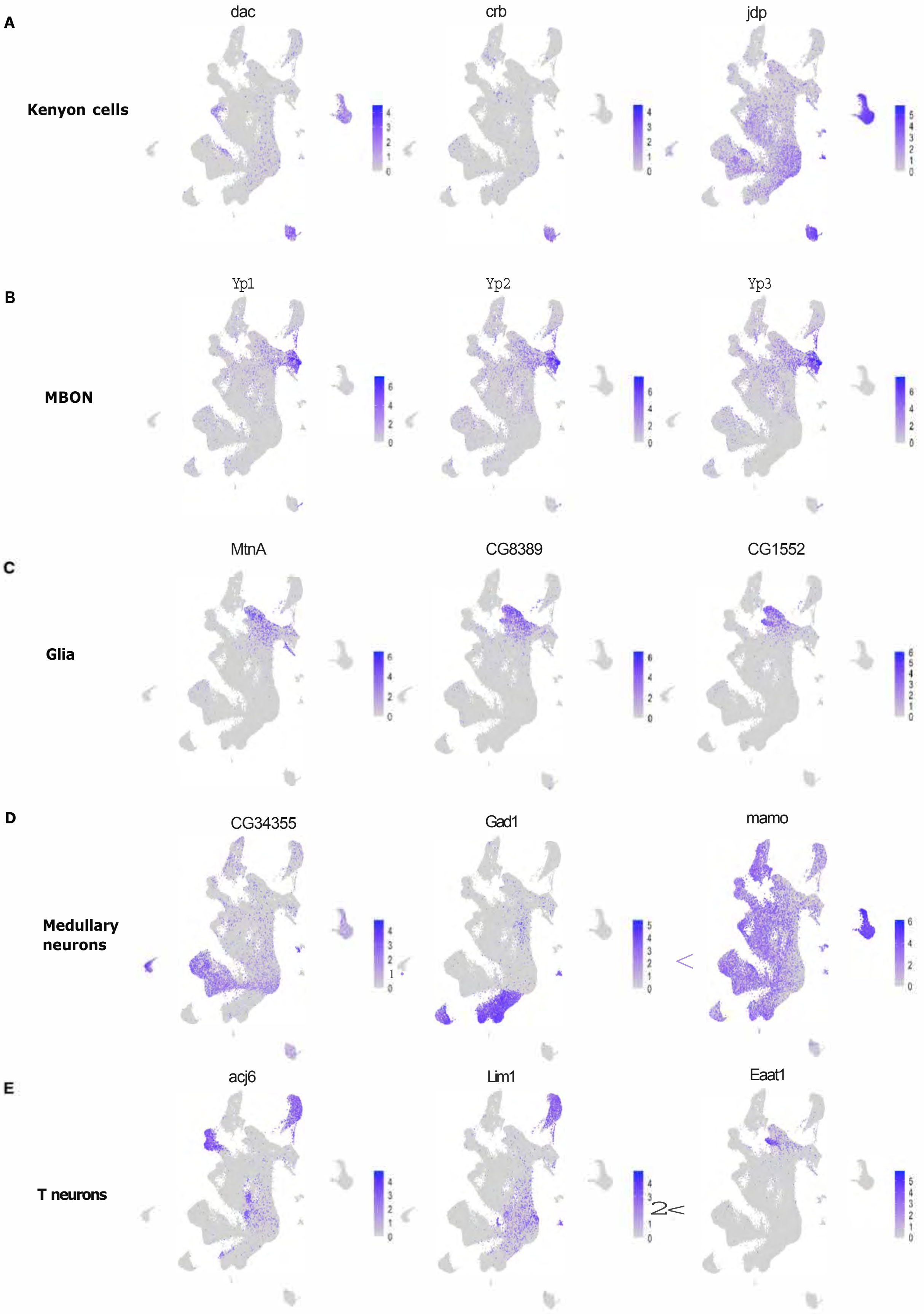
A
Kenyon cells
dac
crb
jdp
B
MBON
Yp1
Yp2
Yp3
C
Glia
MtnA
CG8389
CG1552
D
Medullary neurons
CG34355
Gad1
mamo
E
T neurons
acj6
Lim1
Eaat1

**Supplemental Fig. S5: Individual expression UMAP plots of the top 3 genes used to annotate cellular populations.** Relative expression of the top marker genes, such as dac, crb, and jdp, within the integrated dataset used to annotate Kenyon cells *(A)*. Relative expression of top marker genes, such as Yp1, Yp2, and Yp3, within the integrated dataset used to annotate mushroom body output neurons (MBON) *(B)*. Relative expression of top marker genes, such as MtnA, CG8369, and CG1522, within the integrated dataset annotating the glial cells *(C)*. Relative expression of top marker genes, such as CG34355, Gad1, and mamo, within the integrated dataset, used to annotate the medullary neurons *(D)*. Relative expression of top marker genes, such as acj6, Lim1, and sosie, within the integrated dataset, used to annotate the T neurons *(E)*.

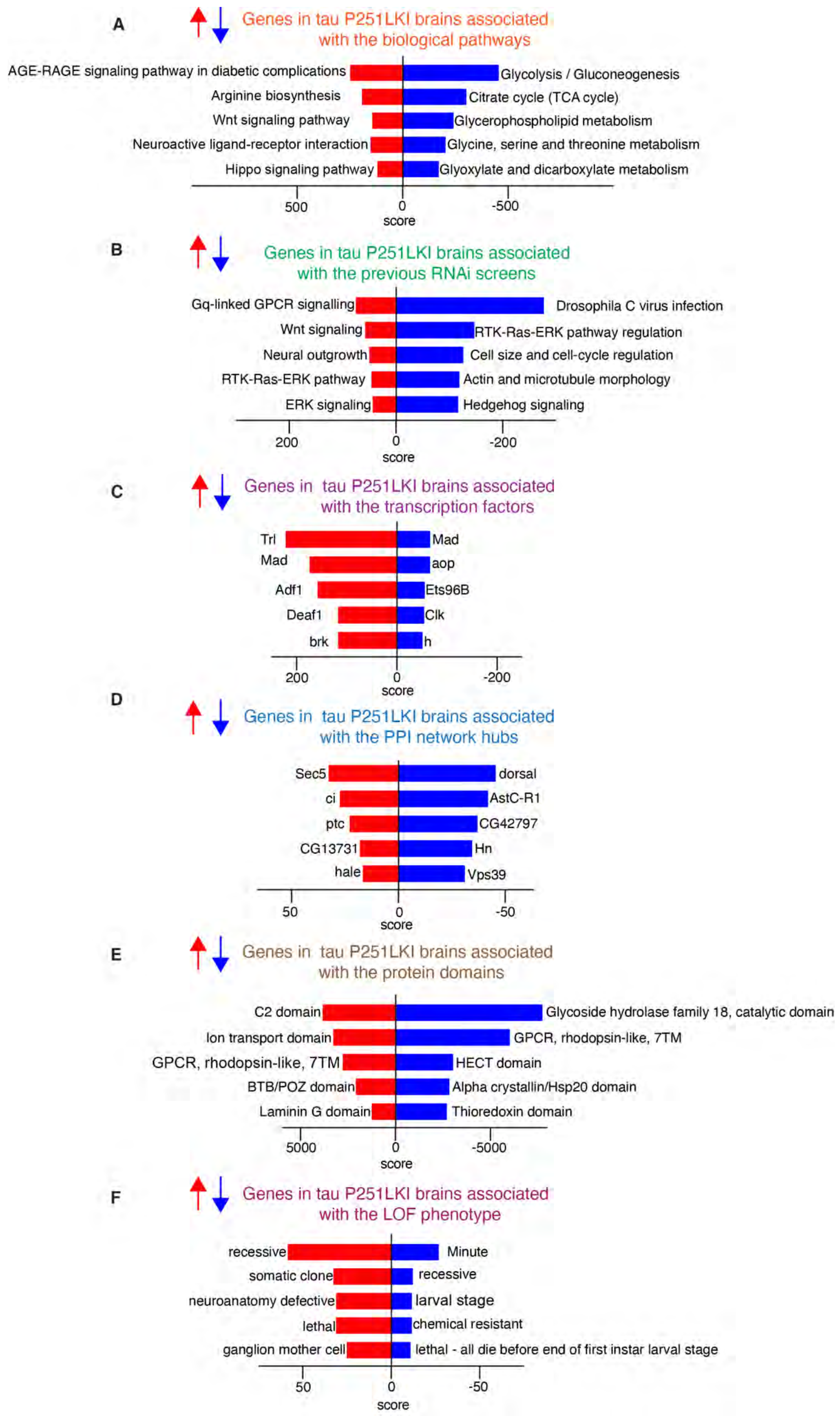
A
Genes in tau P251LKI brains associated with the biological pathways
AGE-RAGE signaling pathway in diabetic complications
Arginine biosynthesis
Wnt signaling pathway
Neuroactive ligand-receptor interaction
Hippo signaling pathway
Glycolysis / Gluconeogenesis
Citrate cycle (TCA cycle)
Glycerophospholipid metabolism
Glycine, serine and threonine metabolism
Glyoxylate and dicarboxylate metabolism
500
0
-500
score
B
Genes in tau P251LKI brains associated with the previous RNAi screens
Gq-linked GPCR signalling
Wnt signaling
Neural outgrowth
RTK-Ras-ERK pathway
ERK signaling
Drosophila C virus infection
RTK-Ras-ERK pathway regulation
Cell size and cell-cycle regulation
Actin and microtubule morphology
Hedgehog signaling
200
0
-200
score
C
Genes in tau P251LKI brains associated with the transcription factors
Trl
Mad
Adf1
Deaf1
brk
Mad
aop
Ets96B
Clk
h
200
0
-200
score
D
Genes in tau P251LKI brains associated with the PPI network hubs
Sec5
ci
ptc
CG13731
hale
dorsal
AstC-R1
CG42797
Hn
Vps39
50
0
-50
score
E
Genes in tau P251LKI brains associated with the protein domains
C2 domain
Ion transport domain
GPCR, rhodopsin-like, 7TM
BTB/POZ domain
Laminin G domain
Glycoside hydrolase family 18, catalytic domain
GPCR, rhodopsin-like, 7TM
HECT domain
Alpha crystallin/Hsp20 domain
Thioredoxin domain
5000
0
-5000
score
F
Genes in tau P251LKI brains associated with the LOF phenotype
recessive
somatic clone
neuroanatomy defective
lethal
ganglion mother cell
Minute
recessive
larval stage
chemical resistant
lethal - all die before end of first instar larval stage
50
0
-50
score

**Supplemental Fig. S6: Differentially regulated enrichment terms in tau P251L KI brains compared to the control.** KEGG pathways analysis of all upregulated genes in tau P251L KI brains identified the AGE-RAGE signaling pathway in diabetic complications and the glycolysis pathway for the downregulated genes in tau P251L KI brains *(A)*. RNAi screen from genome RNAi identified Gq-linked GPCR signaling RNAi screen for the upregulated genes in tau P251L KI brains and *Drosophila* C virus infection screen for the downregulated in tau P251L KI brains *(B)*. The transcription factor analysis identified most of the upregulated genes in tau P251L KI brains associated with Trl transcription factor and downregulated genes in tau P251L KI brains to be associated with Mad transcription factor *(C)*. Protein-protein interactions network hub analysis identified upregulated genes in tau P251L KI brains associated with the Sec5 and downregulated genes in tau P251L KI brains to be associated with dorsal PPI network hub *(D)*. InterPro domain analysis identified the C2 domain as associated with the upregulated genes in tau P251L KI brains and the Glycoside hydrolase family 18 associated with the downregulated genes in tau P251L KI brains *(E)*. Loss of function (LOF) phenotype analysis identified the term recessive associated with the upregulated genes in tau P251L KI brains and minute associated with the downregulated genes in tau P251L KI brains *(F)***.** score = log(p)*z.

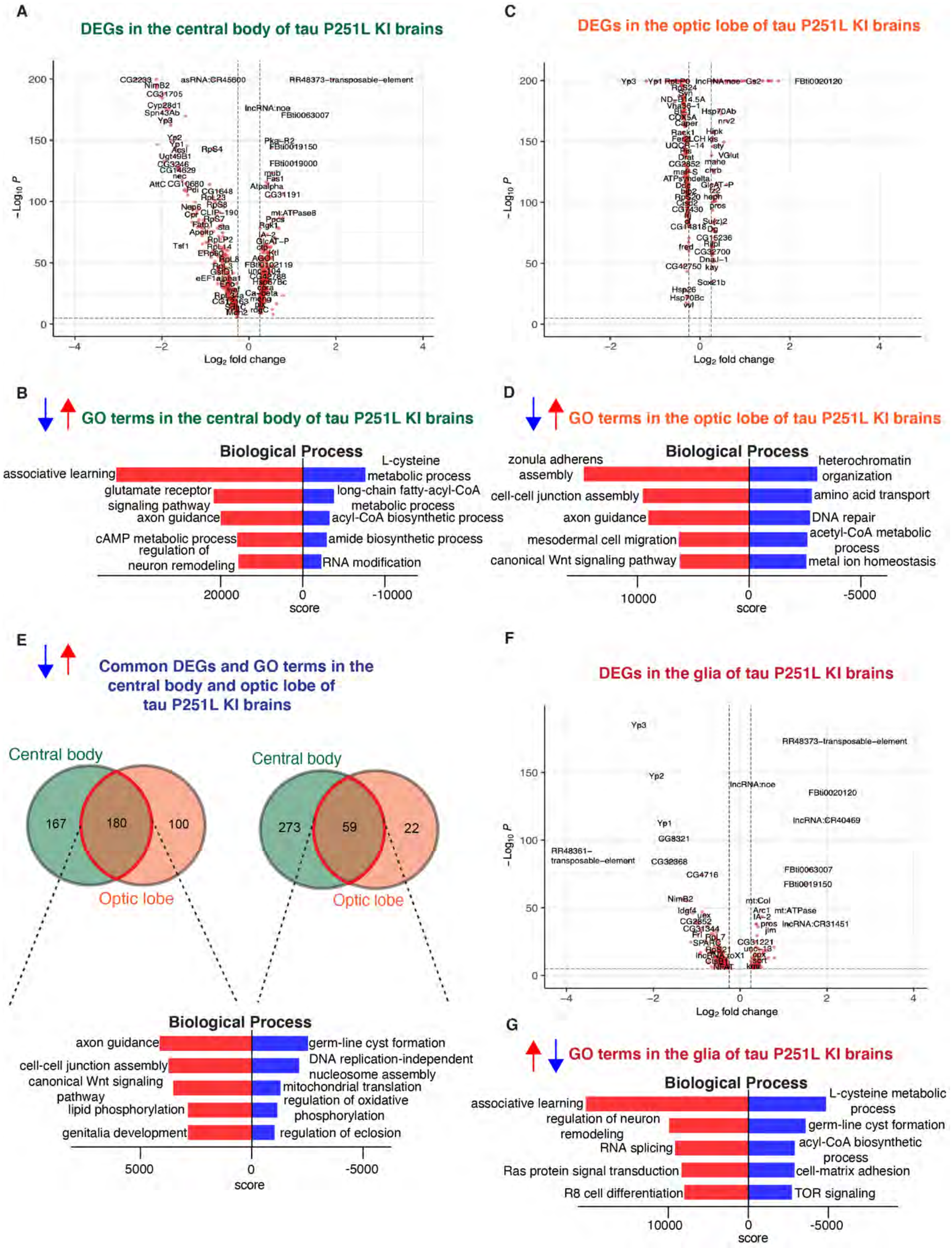

A
DEGs in the central body of tau P251L KI brains
−Log10 P
Log2 fold change
C
DEGs in the optic lobe of tau P251L KI brains
−Log10 P
Log2 fold change
B
GO terms in the central body of tau P251L KI brains
Biological Process
associative learning
glutamate receptor signaling pathway
axon guidance
cAMP metabolic process
regulation of neuron remodeling
L-cysteine metabolic process
long-chain fatty-acyl-CoA metabolic process
acyl-CoA biosynthetic process
amide biosynthetic process
RNA modification
20000
0
-10000
score
D
GO terms in the optic lobe of tau P251L KI brains
Biological Process
zonula adherens assembly
cell-cell junction assembly
axon guidance
mesodermal cell migration
canonical Wnt signaling pathway
heterochromatin organization
amino acid transport
DNA repair
acetyl-CoA metabolic process
metal ion homeostasis
10000
0
-5000
score
E
Common DEGs and GO terms in the central body and optic lobe of tau P251L KI brains
Central body
167
180
100
Optic lobe
Central body
273
59
22
Optic lobe
Biological Process
axon guidance
cell-cell junction assembly
canonical Wnt signaling pathway
lipid phosphorylation
genitalia development
germ-line cyst formation
DNA replication-independent nucleosome assembly
mitochondrial translation
regulation of oxidative phosphorylation
regulation of eclosion
5000
0
-5000
score
F
DEGs in the glia of tau P251L KI brains
−Log10 P
Log2 fold change
G
GO terms in the glia of tau P251L KI brains
Biological Process
associative learning
regulation of neuron remodeling
RNA splicing
Ras protein signal transduction
R8 cell differentiation
L-cysteine metabolic process
germ-line cyst formation
acyl-CoA biosynthetic process
cell-matrix adhesion
TOR signaling
10000
0
-5000
score

**Supplemental Fig. S7: Differential gene expression in the central body, optic lobe, and glia in tau P251L knock-in brains compared to controls.** Differentially regulated genes, both up- and down-regulated, in the central body of tau P251L knock-in brains *(A)*. GO analysis shows biological processes associated with up-regulated and down-regulated genes in the central body of tau P251L knock-in brains *(B)*. Differentially regulated genes, both up and down-regulated, in the optic lobe of tau P251L knock-in brains *(C)*. GO analysis shows biological process associated with up-regulated and down-regulated genes in the optic lobe of tau P251L knock-in brains *(D)*. Biological processes, up and down-regulated, in the common neuronal genes in the central body and optic lobe *(E)*. Differentially regulated genes, both up and down-regulated, in glia of tau P251L knock-in brains *(F)*. GO analysis shows biological process associated with up-regulated and down-regulated genes in glia of tau P251L knock-in brains *(G)*. All dots on the volcano plots are significnat at $FDR < 0.05$ and $log2FC > 0.25$ for up-regulated and $< -0.25$ for down-regulated genes. Score represents the combined score $c = log(p)*z$ (Chen et al. 2013).

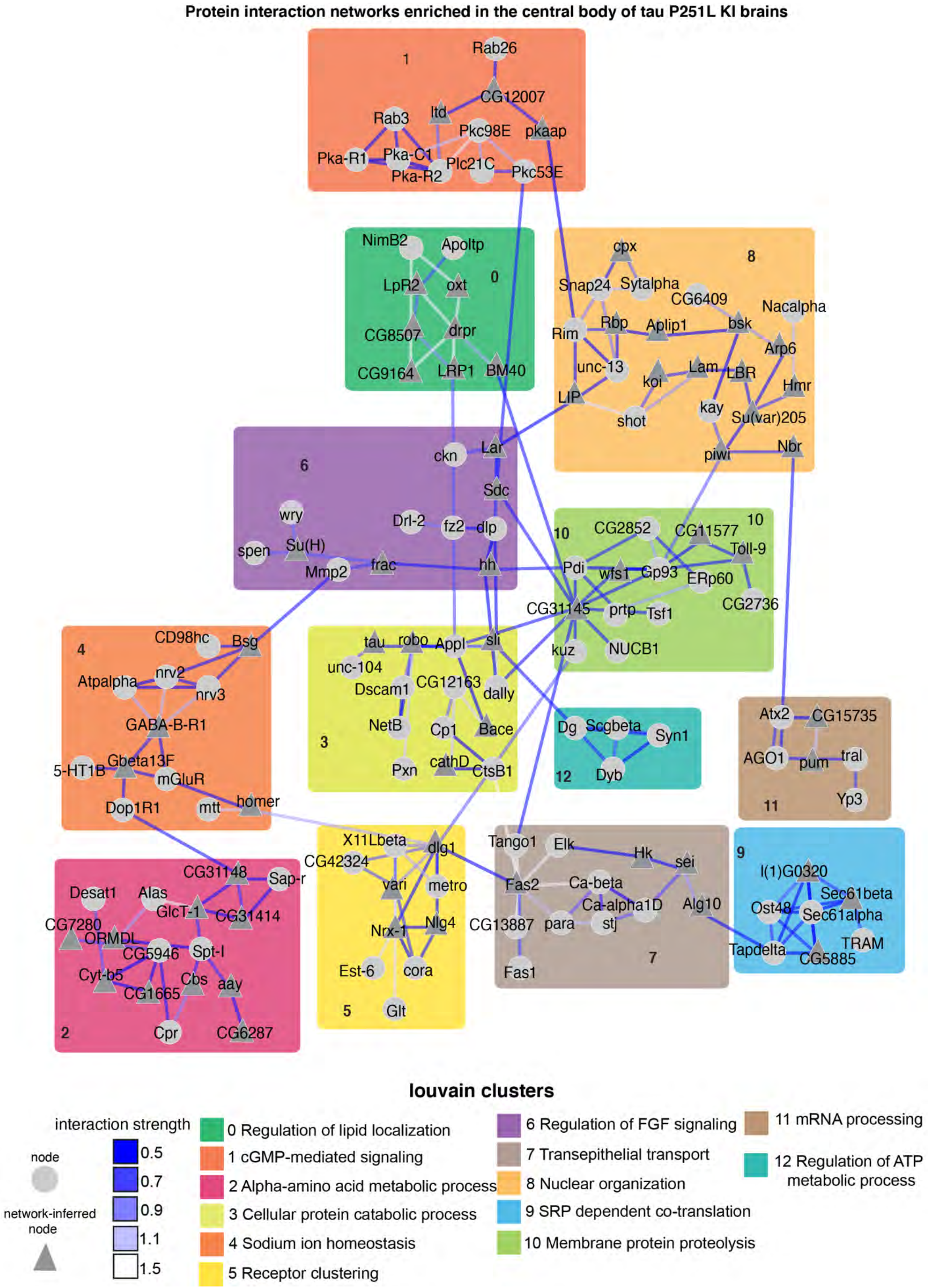

Protein interaction networks enriched in the central body of tau P251L KI brains
louvain clusters
interaction strength
node
network-inferred node
0.5
0.7
0.9
1.1
1.5
0 Regulation of lipid localization
1 cGMP-mediated signaling
2 Alpha-amino acid metabolic process
3 Cellular protein catabolic process
4 Sodium ion homeostasis
5 Receptor clustering
6 Regulation of FGF signaling
7 Transepithelial transport
8 Nuclear organization
9 SRP dependent co-translation
10 Membrane protein proteolysis
11 mRNA processing
12 Regulation of ATP metabolic process

**Supplemental Fig. S8: Protein interaction networks enriched in the central body of tau P251L KI brains.** OmicsIntegrator identified various PPI interaction maps in the central body of the tau P251L KI brain, including regulation of lipid localization, cGMP-mediated signaling, alpha-amino acid transport, cellular protein catabolic process, sodium ion homeostasis, receptor clustering, regulation of FGF signaling, transepithelial transport, nuclear organization, SRP dependent co-translation, membrane protein proteolysis, mRNA processing, and electron transport chain.

**Protein interaction networks enriched in the optic lobe of tau P251L KI brains**

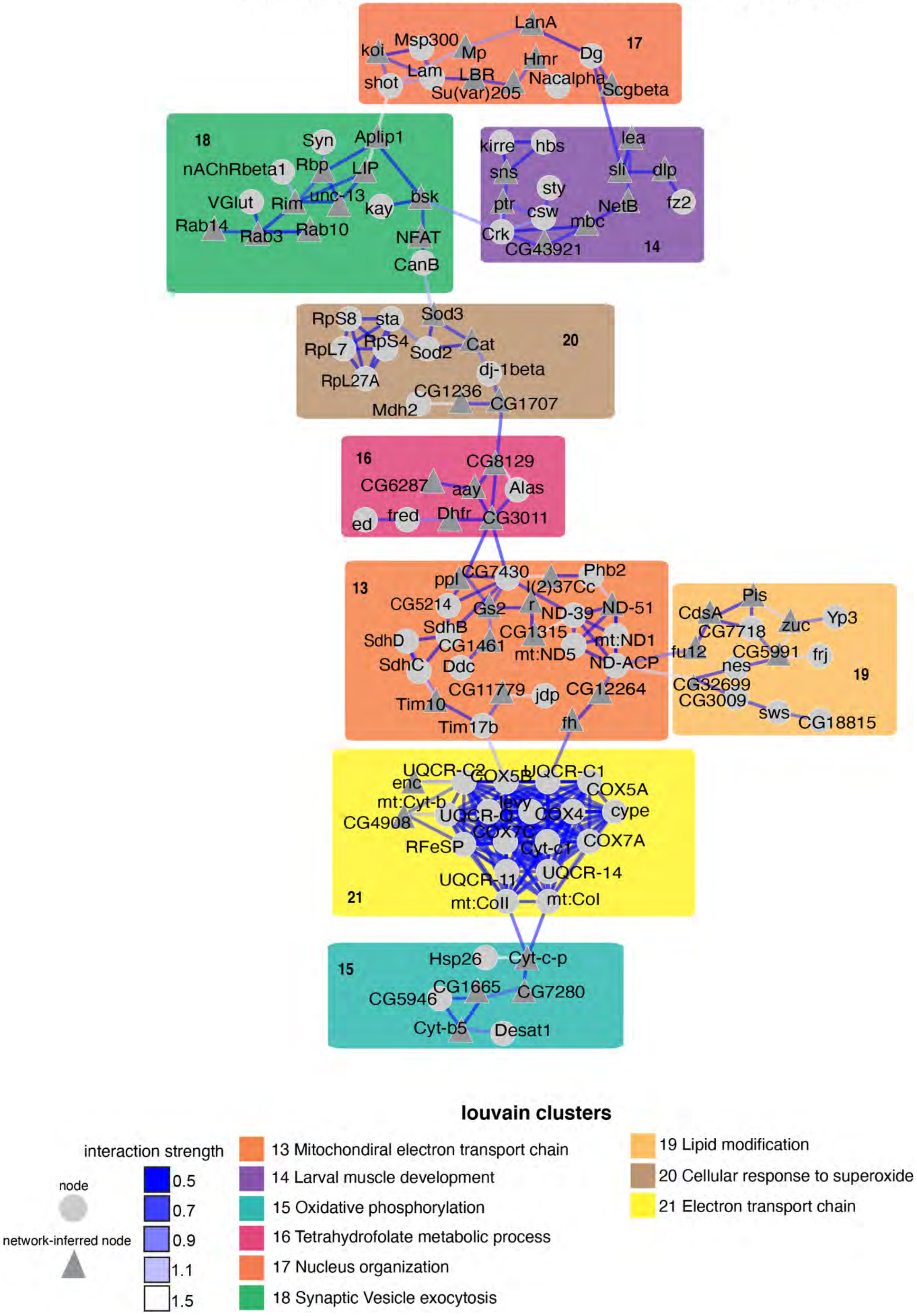

**Supplemental Fig. S9: Protein interaction networks enriched in the optic lobe of tau P251L KI brains.** OmicsIntegrator identified various PPI interaction maps in the optic lobe of the tau P251L KI brain, including mitochondrial electron transport chain, larval muscle development, oxidative phosphorylation, tetrahydrofolate metabolic process, nucleus organization, synaptic vesicle exocytosis, lipid modification, cellular response to superoxide and electron transport chain.

**Protein interaction networks enriched in the glia of tau P251L KI brains**

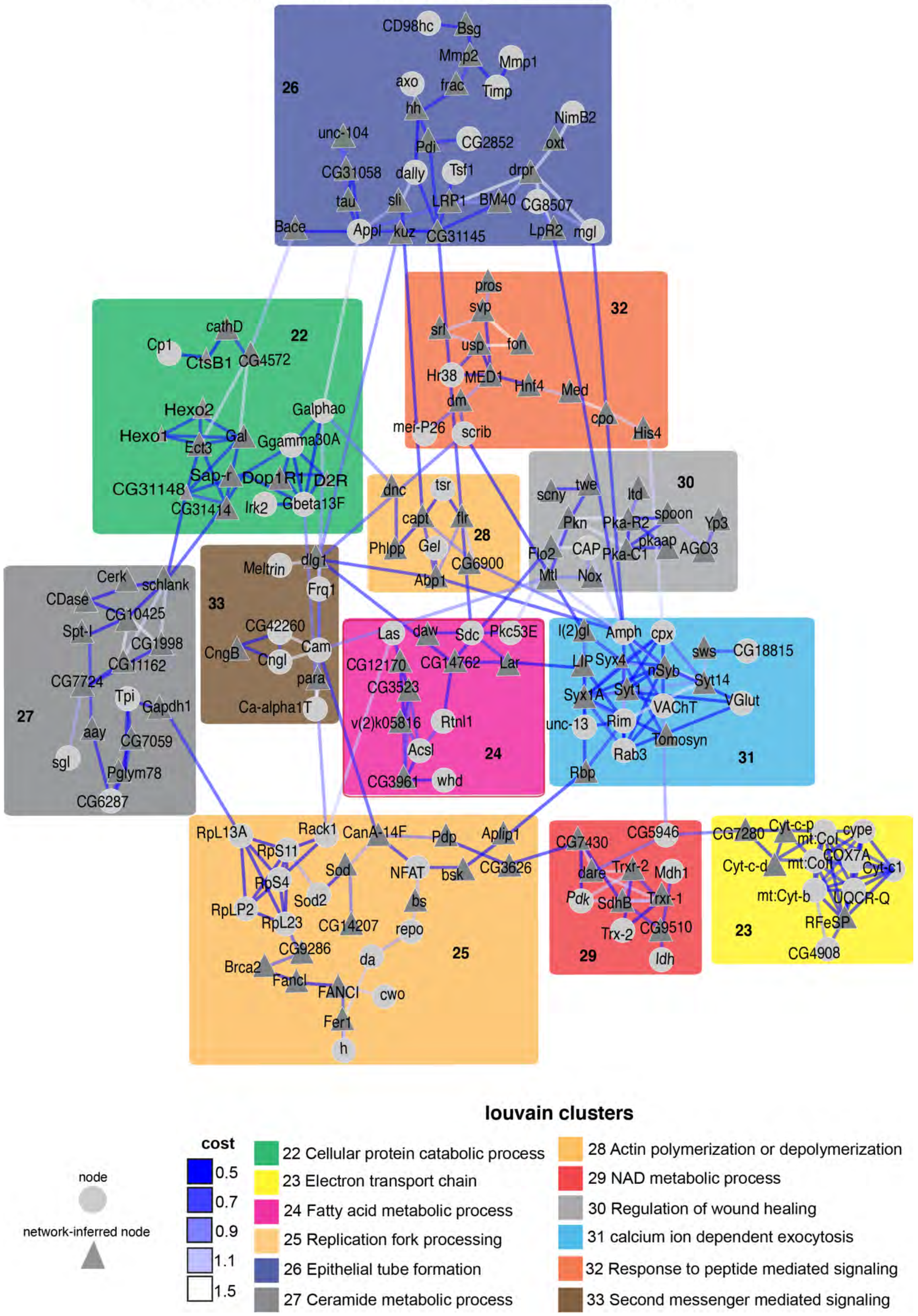

**Supplemental Fig. S10: Protein interaction networks enriched in the glial cells of tau P251L KI brains.** OmicsIntegrator identified various PPI interaction maps in the glial cells of the tau P251L KI brain, including cellular protein catabolic process, electron transport chain, fatty acid metabolic process, replication fork processing, epithelial tube formation, ceramide metabolic process, actin polymerization or depolymerization, NAD metabolic process, regulation of wound healing, calcium ion-dependent exocytosis, response to peptide-mediated signaling, second messenger mediated signaling.

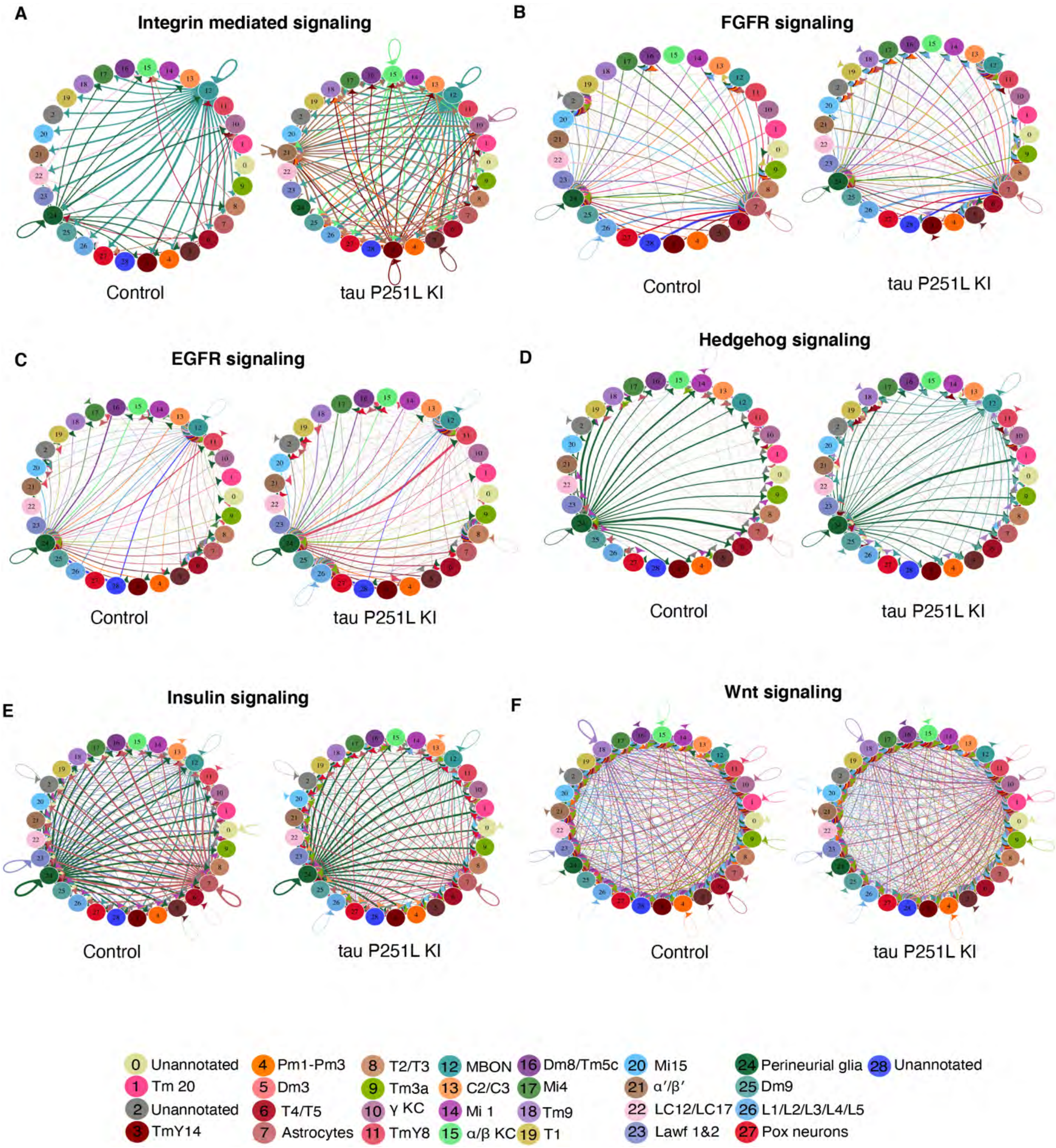
A
Integrin mediated signaling
Control
tau P251L KI
B
FGFR signaling
Control
tau P251L KI
C
EGFR signaling
Control
tau P251L KI
D
Hedgehog signaling
Control
tau P251L KI
E
Insulin signaling
Control
tau P251L KI
F
Wnt signaling
Control
tau P251L KI
0 Unannotated
1 Tm 20
2 Unannotated
3 TmY14
4 Pm1-Pm3
5 Dm3
6 T4/T5
7 Astrocytes
8 T2/T3
9 Tm3a
10 γ KC
11 TmY8
12 MBON
13 C2/C3
14 Mi 1
15 α/β KC
16 Dm8/Tm5c
17 Mi4
18 Tm9
19 T1
20 Mi15
21 α′/β′
22 LC12/LC17
23 Lawf 1&2
24 Perineurial glia
25 Dm9
26 L1/L2/L3/L4/L5
27 Pox neurons
28 Unannotated

**Supplemental Fig. S11: Compared to the control, the cell-cell communication analysis of tau P251L KI brain reveals a distinctive role of glial and Kenyon cells in tau P251L KI brains**. Perineurial glial cells actively send ligands of integrin-mediated signaling to other clusters in the control brain, but perineurial don't send these signals in tau P251L KI brains *(A)*. EGFR signaling increases from perineurial glial cells to the other tau P251L KI brain clusters. Further, 4 clusters send ligands of the EGFR signaling in control, but 5 clusters send ligands in tau P251L KI brains *(B)*. Like EGFR signaling, FGFR signaling increases in tau P251L KI brains: 4 clusters send ligands in the control brain, but 8 clusters send ligands in tau P251L KI brains *(C)*. An increase of hedgehog signaling from the perineurial glial cells to other clusters can be observed in tau P251L KI brains. Further, 4 clusters send hedgehog signaling ligands in the control brains, but 5 clusters in tau P251L KI brains *(D).* A decrease of insulin (10 clusters send ligands in the control while 8 clusters send ligands in tau P251L KI brains) and Wnt signaling (decreased communication lines from cluster 18 in tau P251L KI brains) in tau P251L KI brains *(E&F)***.**

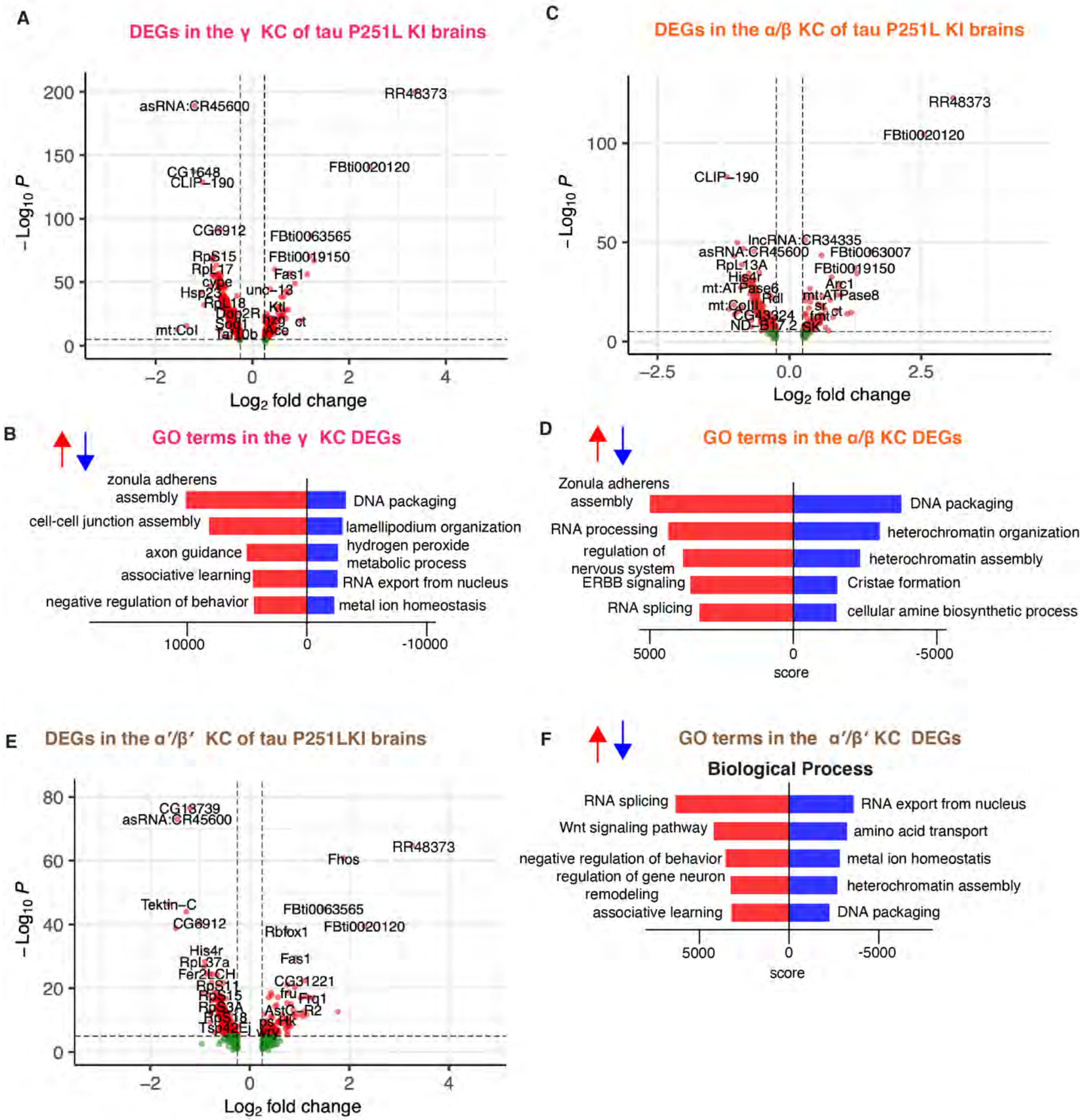
A
DEGs in the γ KC of tau P251L KI brains
RR48373
asRNA:CR45600
CG1648
CLIP-190
FBti0020120
CG6912
FBti0063565
RpS15
FBti0019150
RpL17
Fas1
cype
Hsp23
unc-13
RpL18
Dop2R
Ktl
Scp1
Ace
ct
mt:CoI
Tario0b
−Log10 P
Log2 fold change
B
GO terms in the γ KC DEGs
zonula adherens assembly
cell-cell junction assembly
axon guidance
associative learning
negative regulation of behavior
DNA packaging
lamellipodium organization
hydrogen peroxide metabolic process
RNA export from nucleus
metal ion homeostasis
10000
0
-10000
C
DEGs in the α/β KC of tau P251L KI brains
RR48373
FBti0020120
CLIP-190
lncRNA:CR34335
asRNA:CR45600
FBti0063007
RpL13A
FBti0019150
His4r
Arc1
mt:ATPase6
mt:ATPase8
Rdl
mt:CoIII
sr
CG43324
fmt
ct
ND-B17.2
SK
−Log10 P
Log2 fold change
D
GO terms in the α/β KC DEGs
Zonula adherens assembly
RNA processing
regulation of nervous system
ERBB signaling
RNA splicing
DNA packaging
heterochromatin organization
heterochromatin assembly
Cristae formation
cellular amine biosynthetic process
5000
0
-5000
score
E
DEGs in the α′/β′ KC of tau P251LKI brains
CG13739
asRNA:CR45600
RR48373
Fhos
Tektin-C
FBti0063565
CG6912
FBti0020120
Rbfox1
His4r
RpL37a
Fas1
Fer2LCH
RpS11
CG31221
RpS15
fru
Frq1
RpS3A
AstC-R2
RpS18
Hk
Tsp42Ej
−Log10 P
Log2 fold change
F
GO terms in the α′/β′ KC DEGs
Biological Process
RNA splicing
Wnt signaling pathway
negative regulation of behavior
regulation of gene neuron remodeling
associative learning
RNA export from nucleus
amino acid transport
metal ion homeostatis
heterochromatin assembly
DNA packaging
5000
0
-5000
score

**Supplemental Fig. S12: Differential gene expression and enrichment analysis in the Kenyon cells (KC) of tau P251L KI brains compared to the control.** The volcano plot shows differentially up and down-regulated genes in the $\gamma$ KC cluster *(A)*. Gene ontology analysis found zonula adherens assembly to be the most common upregulated and DNA packaging to be the most common downregulated biological process in the $\gamma$ KC cluster *(B)* The volcano plot shows differentially up and down-regulated genes in the $\alpha/\beta$ KC cluster *(C)*. Gene ontology analysis found zonula adherens assembly to be the most common upregulated and DNA packaging to be the most common downregulated biological process in the $\alpha/\beta$ KC cluster *(D)*. The volcano plot shows differentially up and down-regulated genes in the $\alpha'/\beta'$ KC cluster *(F)*. Gene ontology analysis found RNA splicing to be the most common upregulated and RNA export from the nucleus to be the most common downregulated biological process in the $\alpha'/\beta'$ KC cluster *(F)*. All red dots on the volcano plots are significant genes meeting the cutoff of FDR-adjusted p-value < 0.05 and log2FC > 0.25 for upregulated and < - 0.25 for downregulated genes, score = log(p)*z.

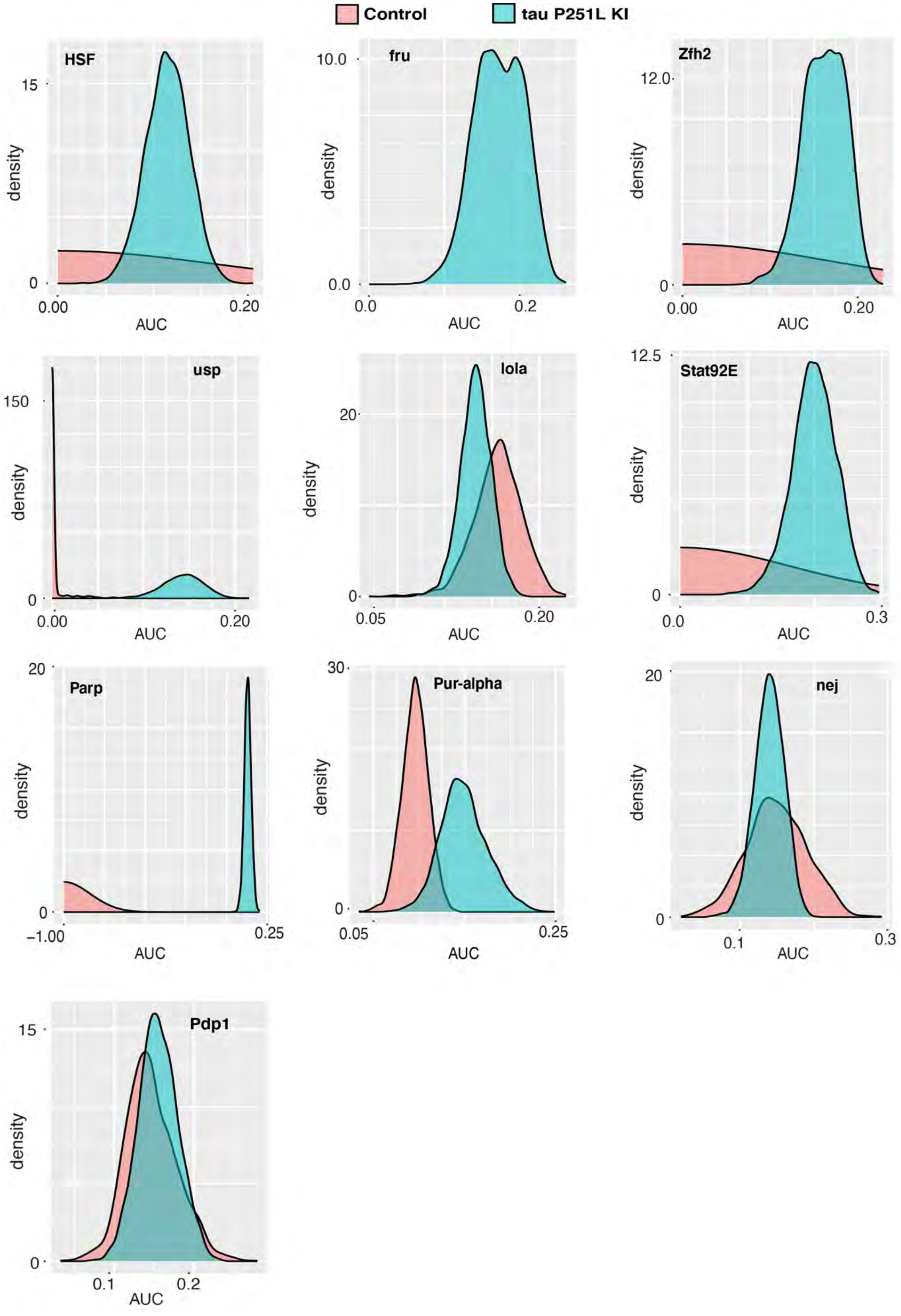

Control
tau P251L KI
HSF
fru
Zfh2
usp
lola
Stat92E
Parp
Pur-alpha
nej
Pdp1
density
AUC

**Supplemental Fig. S13: Density plots compare the expression of regulons enriched in the tau P251L knock-in Kenyon cells compared to controls.** The density plots comparing the expression of regulons in the control vs tau P251L knock-in Kenyon cells show that the expression of regulons such as HSF, fru, Zfh2, usp, Stat92E, and Parp is significantly elevated in the tau P251L knock-in Kenyon cells compared to the control.
