## Supplemental Code for "Transcriptional programs mediating neuronal toxicity and altered glial-neuronal signaling in a *Drosophila* knock-in tauopathy model"

**Supplemental code**
**The following code was used to generate Supplemental Figure S3**

```
#First load the libraries

library(dplyr)
library(Seurat)
library(patchwork)
library(ggplot2)
library(stringr)
library(SoupX)
library(reticulate)
library(glmpca)
library(SeuratWrappers)
library(scry)
library(reticulate)
library(monocle3)
library(FlexDotPlot)
library(cowplot)
library(tidyverse)
library(viridis)
library(scCustomize)
library(qs)
library(gridExtra)
library(plyr)
library(circlize)
library(ComplexHeatmap)
library(EnhancedVolcano)
library(data.table)
library(magrittr)
library(raster)
library(rgdal)
library(classInt)
library(RColorBrewer)
library(Seurat)
library(SeuratDisk)

rm(scrna.combined.seurat.hassan072022)

#load the object
scrna.combined.seurat.hassan072022 <- readRDS("/Volumes/Macintosh
HD/Users/hassanbukhari/Desktop/OneDrive - Mass General Brigham/Tau P251LKI figures
updated-02-27-223/New Seurat Object-2023/seurat_multi_rerun_annotations
2/scrna.combined.seurat.hassan_2022.rds")


p1 <- DimPlot(scrna.combined.seurat.hassan072022, reduction = "umap", label = TRUE, raster
= FALSE, repel = TRUE)
p1
```

```
#Rename the Seurat Object to scrna.combined

scrna.combined <-  scrna.combined.seurat.hassan072022
rm(scrna.combined.seurat.hassan072022)

#Now subset all the control. Save them and count their cells.


# Split the control1 object
Control1 <- subset(scrna.combined, subset= orig.ident=="control1")
p1 <- DimPlot(Control1, reduction = "umap", label = FALSE, repel = TRUE)
p1

saveRDS(Control1, paste0("/Volumes/Macintosh HD/Users/hassanbukhari/Desktop/OneDrive - Mass General Brigham/Tau P251LKI figures updated-02-27-223/Figure 3/Figure 3 Supplementary files/", "Control1.rds"))

md1<- Control1@meta.data %>% as.data.table

a <-  md1[, .N, by = c("seurat_clusters")]

write.table (a, paste0("/Volumes/Macintosh HD/Users/hassanbukhari/Desktop/OneDrive - Mass General Brigham/Tau P251LKI figures updated-02-27-223/Figure 3/Figure 3 Supplementary files/", "Control1-Table.xls"),sep='\t',row.names=F,col.names=T, quote=F)

# Split the control3 object
Control3 <- subset(scrna.combined, subset= orig.ident=="CRN00224919")
p2 <- DimPlot(Control3, reduction = "umap", label = FALSE, repel = TRUE)
p2

saveRDS(Control3, paste0("/Volumes/Macintosh HD/Users/hassanbukhari/Desktop/OneDrive - Mass General Brigham/Tau P251LKI figures updated-02-27-223/Figure 3/Figure 3 Supplementary files/", "Control3.rds"))

md1<- Control3@meta.data %>% as.data.table

a <-  md1[, .N, by = c("seurat_clusters")]

write.table (a, paste0("/Volumes/Macintosh HD/Users/hassanbukhari/Desktop/OneDrive - Mass General Brigham/Tau P251LKI figures updated-02-27-223/Figure 3/Figure 3 Supplementary files/", "Control3-Table.xls"),sep='\t',row.names=F,col.names=T, quote=F)


# Split the control2 object
Control2 <- subset(scrna.combined, subset= orig.ident=="CTRL2merged")
p2 <- DimPlot(Control2, reduction = "umap", label = FALSE, repel = TRUE)
p2

saveRDS(Control2, paste0("/Volumes/Macintosh HD/Users/hassanbukhari/Desktop/OneDrive - Mass General Brigham/Tau P251LKI figures updated-02-27-223/Figure 3/Figure 3 Supplementary files/", "Control2.rds"))
```

```
md1<- Control2@meta.data %>% as.data.table

a <-  md1[, .N, by = c("seurat_clusters")]

write.table (a, paste0("/Volumes/Macintosh HD/Users/hassanbukhari/Desktop/OneDrive - Mass
General Brigham/Tau P251LKI figures updated-02-27-223/Figure 3/Figure 3 Supplementary
files/", "Control2-Table.xls"),sep='\t',row.names=F,col.names=T, quote=F)

# Split the P251L1 object
P251L1 <- subset(scrna.combined, subset= orig.ident=="P251L1merged")
p1 <- DimPlot(P251L1, reduction = "umap", label = FALSE, repel = TRUE)
p1

saveRDS(P251L1, paste0("/Volumes/Macintosh HD/Users/hassanbukhari/Desktop/OneDrive -
Mass General Brigham/Tau P251LKI figures updated-02-27-223/Figure 3/Figure 3
Supplementary files/", "P251L1.rds"))

md1<- P251L1@meta.data %>% as.data.table

a <-  md1[, .N, by = c("seurat_clusters")]

write.table (a, paste0("/Volumes/Macintosh HD/Users/hassanbukhari/Desktop/OneDrive - Mass
General Brigham/Tau P251LKI figures updated-02-27-223/Figure 3/Figure 3 Supplementary
files/", "P251L1.xls"),sep='\t',row.names=F,col.names=T, quote=F)

# Split the P251L2 object
P251L2 <- subset(scrna.combined, subset= orig.ident=="P251L2merged")
p1 <- DimPlot(P251L2, reduction = "umap", label = FALSE, repel = TRUE)
p1

saveRDS(P251L2, paste0("/Volumes/Macintosh HD/Users/hassanbukhari/Desktop/OneDrive -
Mass General Brigham/Tau P251LKI figures updated-02-27-223/Figure 3/Figure 3
Supplementary files/", "P251L2.rds"))

md1<- P251L2@meta.data %>% as.data.table

a <-  md1[, .N, by = c("seurat_clusters")]

write.table (a, paste0("/Volumes/Macintosh HD/Users/hassanbukhari/Desktop/OneDrive - Mass
General Brigham/Tau P251LKI figures updated-02-27-223/Figure 3/Figure 3 Supplementary
files/", "P251L2.xls"),sep='\t',row.names=F,col.names=T, quote=F)


# Split the P251L3 object
P251L3 <- subset(scrna.combined, subset= orig.ident=="LIB055588_CRN00233457")
p1 <- DimPlot(P251L3, reduction = "umap", label = FALSE, repel = TRUE)
p1
```

```
saveRDS(P251L3, paste0("/Volumes/Macintosh HD/Users/hassanbukhari/Desktop/OneDrive -
Mass General Brigham/Tau P251LKI figures updated-02-27-223/Figure 3/Figure 3
Supplementary files/", "P251L3.rds"))

md1<- P251L3@meta.data %>% as.data.table

a <-  md1[, .N, by = c("seurat_clusters")]

write.table (a, paste0("/Volumes/Macintosh HD/Users/hassanbukhari/Desktop/OneDrive - Mass
General Brigham/Tau P251LKI figures updated-02-27-223/Figure 3/Figure 3 Supplementary
files/", "P251L3.xls"),sep='\t',row.names=F,col.names=T, quote=F)


#Split the controls together
Controls <- subset(scrna.combined, subset= treatment =="control")

p1 <- DimPlot(Controls, reduction = "umap", label = FALSE, repel = FALSE)
p1

saveRDS(Controls, paste0("/Volumes/Macintosh HD/Users/hassanbukhari/Desktop/OneDrive -
Mass General Brigham/Tau P251LKI figures updated-02-27-223/Figure 3/Figure 3
Supplementary files/", "Controls.rds"))

md1<- Controls@meta.data %>% as.data.table

a <-  md1[, .N, by = c("seurat_clusters")]

write.table (a, paste0("/Volumes/Macintosh HD/Users/hassanbukhari/Desktop/OneDrive - Mass
General Brigham/Tau P251LKI figures updated-02-27-223/Figure 3/Figure 3 Supplementary
files//", "Controls.xls"),sep='\t',row.names=F,col.names=T, quote=F)

#Split the Knock ins together

rm(scrna.combined.seurat.hassan072022)

P251Ls <- subset(scrna.combined, subset= treatment =="P251L")


#load the knock in

#P251Ls  <- readRDS(paste0("/Volumes/shb27/SC-Hassan/Seurat Multi3/Samples
separated/P251Ls/P251Ls.rds"))

p1 <- DimPlot(P251Ls, reduction = "umap", label = FALSE, repel = TRUE)
p1

saveRDS(P251Ls, paste0("/Volumes/Macintosh HD/Users/hassanbukhari/Desktop/OneDrive -
Mass General Brigham/Tau P251LKI figures updated-02-27-223/Figure 3/Figure 3
Supplementary files/", "P251Ls.rds"))

md1<- P251Ls@meta.data %>% as.data.table
```

```
a <-  md1[, .N, by = c("seurat_clusters")]

write.table (a, paste0("/Volumes/Macintosh HD/Users/hassanbukhari/Desktop/OneDrive - Mass General Brigham/Tau P251LKI figures updated-02-27-223/Figure 3/Figure 3 Supplementary files/", "P251Ls.xls"),sep='\t',row.names=F,col.names=T, quote=F)


#to show representative Cholinergic

FeaturePlot(scrna.combined.seurat.hassan072022, "FBgn0270928", raster = FALSE) +
scale_colour_gradientn(colours = brewer.pal(n = 9, name = "PuBu"))

#to show representative GABAergic

FeaturePlot(scrna.combined.seurat.hassan072022, "FBgn0004516", raster = FALSE) +
scale_colour_gradientn(colours = brewer.pal(n = 9, name = "Greys"))

#to show representative Glutamatergic

FeaturePlot(scrna.combined.seurat.hassan072022, "FBgn0031424", raster = FALSE) +
scale_colour_gradientn(colours = brewer.pal(n = 9, name = "Greens"))

#to show representative Dopaminergic

FeaturePlot(scrna.combined.seurat.hassan072022, "FBgn0034136", raster = FALSE) +
scale_colour_gradientn(colours = brewer.pal(n = 9, name = "BuPu"))



#Now to determine the cells expressing > 2 of the gene within the all the datasets


#For Control 1

Control1
Control1 <- subset(scrna.combined, subset= orig.ident=="control1")

#cholinergic.neurons
cholinergic.neurons <- subset(x = Control1, subset = FBgn0270928 > 2)
cholinergic.neurons

#Glutamatergic.neurons
Glutamatergic.neurons <- subset(x = Control1, subset = FBgn0031424 > 2)
Glutamatergic.neurons

#GABAergic.neurons
GABAergic.neurons <- subset(x = Control1, subset = FBgn0004516 > 2)
GABAergic.neurons

#Dopaminergic.neurons
```

```
Dopaminergic.neurons <- subset(x = Control1, subset = FBgn0034136 > 2)
Dopaminergic.neurons


#For Control2

Control2 <- subset(scrna.combined, subset= orig.ident=="CTRL2merged")
Control2
#cholinergic.neurons
cholinergic.neurons <- subset(x = Control2, subset = FBgn0270928 > 2)
cholinergic.neurons

#Glutamatergic.neurons
Glutamatergic.neurons <- subset(x = Control2, subset = FBgn0031424 > 2)
Glutamatergic.neurons

#GABAergic.neurons
GABAergic.neurons <- subset(x = Control2, subset = FBgn0004516 > 2)
GABAergic.neurons

#Dopaminergic.neurons
Dopaminergic.neurons <- subset(x = Control2, subset = FBgn0034136 > 2)
Dopaminergic.neurons

# For Control3
Control3 <- subset(scrna.combined, subset= orig.ident=="CRN00224919")
Control3
#cholinergic.neurons
cholinergic.neurons <- subset(x = Control3, subset = FBgn0270928 > 2)
cholinergic.neurons

#Glutamatergic.neurons
Glutamatergic.neurons <- subset(x = Control3, subset = FBgn0031424 > 2)
Glutamatergic.neurons

#GABAergic.neurons
GABAergic.neurons <- subset(x = Control3, subset = FBgn0004516 > 2)
GABAergic.neurons

#Dopaminergic.neurons
Dopaminergic.neurons <- subset(x = Control3, subset = FBgn0034136 > 2)
Dopaminergic.neurons

#For P251L1
P251L1 <- subset(scrna.combined, subset= orig.ident=="P251L1merged")
P251L1

#cholinergic.neurons
cholinergic.neurons <- subset(x = P251L1, subset = FBgn0270928 > 2)
cholinergic.neurons
```

```
#Glutamatergic.neurons
Glutamatergic.neurons <- subset(x = P251L1, subset = FBgn0031424 > 2)
Glutamatergic.neurons

#GABAergic.neurons
GABAergic.neurons <- subset(x = P251L1, subset = FBgn0004516 > 2)
GABAergic.neurons

#Dopaminergic.neurons
Dopaminergic.neurons <- subset(x = P251L1, subset = FBgn0034136 > 2)
Dopaminergic.neurons


#For P251L2
P251L2 <- subset(scrna.combined, subset= orig.ident=="P251L2merged")
P251L2

#cholinergic.neurons
cholinergic.neurons <- subset(x = P251L2, subset = FBgn0270928 > 2)
cholinergic.neurons

#Glutamatergic.neurons
Glutamatergic.neurons <- subset(x = P251L2, subset = FBgn0031424 > 2)
Glutamatergic.neurons

#GABAergic.neurons
GABAergic.neurons <- subset(x = P251L2, subset = FBgn0004516 > 2)
GABAergic.neurons

#Dopaminergic.neurons
Dopaminergic.neurons <- subset(x = P251L2, subset = FBgn0034136 > 2)
Dopaminergic.neurons


#For P251L3
P251L3 <- subset(scrna.combined, subset= orig.ident=="LIB055588_CRN00233457")
P251L3

#cholinergic.neurons
cholinergic.neurons <- subset(x = P251L3, subset = FBgn0270928 > 2)
cholinergic.neurons

#Glutamatergic.neurons
Glutamatergic.neurons <- subset(x = P251L3, subset = FBgn0031424 > 2)
Glutamatergic.neurons

#GABAergic.neurons
GABAergic.neurons <- subset(x = P251L3, subset = FBgn0004516 > 2)
GABAergic.neurons

#Dopaminergic.neurons
```

```
Dopaminergic.neurons <- subset(x = P251L3, subset = FBgn0034136 > 2)
Dopaminergic.neurons


#For the feature plots use the following code

#Kenyon cells #jdp #dac #crb
FeaturePlot(scrna.combined, features = c("FBgn0027654", "FBgn0005677", "FBgn0259685"),
raster = FALSE)


#MBON #Yp1 #Yp2 #Yp3
FeaturePlot(scrna.combined, features = c("FBgn0004045", "FBgn0005391", "FBgn0004047"),
raster = FALSE)

#Glia #MtnA #CG8369 #CG1552
FeaturePlot(scrna.combined, features = c("FBgn0002868", "FBgn0040532", "FBgn0030258"),
raster = FALSE)

#Medullary neurons #CG34355 #Gad1 #mamo
FeaturePlot(scrna.combined, features = c("FBgn0085384", "FBgn0004516", "FBgn0267033"),
raster = FALSE)

# T neurons #acj6 #Lim1 #Eaat1
FeaturePlot(scrna.combined, features = c("FBgn0000028", "FBgn0026411", "FBgn0026439"),
raster = FALSE)
```
