## Supplemental Methods for "Transcriptional programs mediating neuronal toxicity and altered glial-neuronal signaling in a *Drosophila* knock-in tauopathy model"

### Sectioning, immunostaining and imaging

Adult flies were fixed in formalin at 1, 10 and 30 days of age and embedded in paraffin. Serial frontal sections including the entire brain were prepared. Sections were stained with hematoxylin and eosin to assess brain vacuolization, which was quantified by counting vacuoles larger than 5 microns throughout the entire brain. For immunostaining, antigen retrieval was performed by microwaving the sections in 10 mM sodium citrate buffer. Immunohistochemistry was performed with the avidin–biotin–peroxidase complex detection method (Vector Laboratories). For immunofluorescence, secondary antibodies coupled to Alexa 488 or Alexa 555 (Invitrogen, 1:200) were used and sections were mounted in DAPI containing mounting media. The number of PCNA-positive cells throughout the entire brain was counted following immunostaining. For quantification of pH2Av, a region of interest comprised of approximately 100 Kenyon neurons was identified in well-oriented sections of the mushroom body and the number of neurons containing one or more than one immuno-positive foci was determined. Images were taken on Zeiss LSM800 confocal microscope (Carl Zeiss, AG), and quantification was performed using Image-J software. For all histological analyses, at least 6 brains were analyzed per genotype and time point. The sample size (n), mean and SEM are given in the figure legends. All statistical analyses were performed using GraphPad Prism 5.0. For comparisons across more than 2 groups, one-way ANOVA with Tukey post-hoc analysis was used. For comparison of 2 groups Student's t-tests were performed.

### Comet assay

Two brains per genotype were dissected from adult flies in ice cold PBS. The brains were homogenized with a plastic pestle and subjected to comet using commercially available reagents (CometAssay, Trevigen). Fifty nuclei were quantified per trail using Casplab software. The experiment was repeated 3 times.

**Measurement of oxygen consumption and extracellular acidification rates**

The OCR and extracellular acidification rate were measured as previously described (Sarkar et al. 2020). Briefly, brains from 10-day-old flies were dissected and plated at one brain per well on XFe96 plates (Seahorse Bioscience) and metabolic parameters were assayed. 6 brains per genotype were analyzed. OCR values were normalized to DNA content using a CyQUANT assay (ThermoFisher) following the manufacturer's protocol.

**scRNA sequencing**

**Sample preparation**

To dissociate fly brains for the scRNA sequencing we modified previously published fly brain dissociation protocols (Li et al. 2017; Davie et al. 2018). Briefly, 20 male and 20 female brains from 10-day-old flies were dissected on ice cold Schneider's medium with FBS (Gibco, filtered 10% FBS). After a brief centrifugation, the supernatant was removed and the brains washed with ice cold PBS to remove Schneider's medium. The brains were then incubated at 25 °C with 300 μl of 0.05% trypsin-EDTA (Fisher Scientific) for 30 minutes, with continuous pipetting every 5 minutes. Additionally, the solution containing brain chunks was passed through 25-gauge needle (25G 5/8), without introducing air bubbles, roughly 50 times. After the brains were fully dissociated, the resultant solution was poured through a 10 μm pluri-select cell strainer (Fisher Scientific) and 400 μl of ice-cold Schneider's medium containing FBS was added to inactivate trypsin. The sample was centrifuged for 15 minutes at 600 x g and the supernatant was removed without disturbing the pellet. The pellet was suspended in sterile PBS containing 0.04% BSA. The cells were quantified, and the viability was determined using AO-PI reagents (Logos Biosystems).

**Single-cell encapsulation, sequencing, and downstream processing**

We proceeded only with samples having more than 90% viability for the single-cell encapsulation. The samples were encapsulated, 6 libraries were prepared, 3 control and 3 tau P251L knock-in, at the single-cell core facility at Harvard Medical School, following the manufacturer's protocol

(10x Genomics). The libraries were sequenced on NovaSeq 6000 V1.5 S2 located in the Harvard Medical School Biopolymers Facility.

**10x raw data processing**

The sequenced libraries were processed using Cellranger (version 6.0, 10x Genomics). The *Drosophila* reference genome BDGP6.32 was used and built following 10x Genomics users guide instruction. The output of Cellranger were used as an input of SoupX (version 1.5.2) and Scrublet (version 0.2.2) to remove potential ambient RNA and doublets, respectively. The resulting count matrices, indicating transcripts (UMI) and cells (barcodes) detected by sequencing, were used for quality control and downstream analysis.

**Seurat data processing**

The ambient RNA and doublet removed count matrices were used as input of Seurat (version 4.1.0). For each library, cells with less than 10% of mitochondrial genes were kept for downstream analysis. In addition, cells with feature counts and UMI counts within 3 standard deviations of the mean library feature counts and UMI counts were kept for downstream processing. In addition, only features detected in at least 3 cells were kept. Data normalization was then performed across libraries to remove variation of sequencing depth, by employing a global-scaling normalization method "LogNormalize" (i.e., the feature expression measurements for each cell were divided by the total expression, multiplied by a scale factor of 10,000, and then log-transformed). The top 2,000 most variable features were selected and used for downstream analysis including dimension reduction. To annotate cell types and perform differential expression analysis between conditions in an experiment, libraries sequenced in one experiment were integrated by identifying common anchors between conditions. The dimensions of the expression matrices were then reduced by principal component analysis (PCA). The ElbowPlot function was used to determine the optimal number of dimensions used to identify cell clusters. Cell clusters were identified with the default method in Seurat. In brief, a K nearest neighbors (KNN) graph was first constructed based on the Euclidean distance of cells in PCA space, and the edge weights between any two cells were refined based on the shared overlap in their local neighborhoods (Jaccard similarity). Louvain algorithm was applied to iteratively group cells together to form optimized communities with the resolution parameter of 0.5. Cells within the graph-based clusters determined above were further visualized and explored by Uniform Manifold Approximation and Projection (UMAP) non-linear dimensional reduction (Becht et al. 2018). The cell clusters were then annotated based on manual inspection of top feature genes and known marker genes expression in each cluster. Finally, the differentially expressed genes across conditions were examined by comparing expression profile of each cluster across experimental conditions using MAST 1.23.1 (Finak et al. 2015). The significant differentially expressed genes (DEGs) were defined as FDR-adjusted p-value < 0.05 and absolute log2fold-change > 0.25. The top DEGs were also shown in the volcano plots and the heatmap.

**Cell cluster annotation, gene enrichment and ontology analyses**

To annotate cluster identity, we used data collated by the *Drosophila* RNAi Screening Center single-cell RNA sequencing DataBase (DRscDB) from previous *Drosophila* brain single cell sequencing analyses (Davie et al. 2018; Li et al. 2022; Hu et al. 2021). Gene enrichment and ontology analyses were performed with FlyEnrichr (Chen et al. 2013; Kuleshov et al. 2016) using 2 differentially enriched gene lists: 1) differentially up-regulated genes, FDR-adjusted p-value < 0.05 and log2fold-change > 0.25, throughout all brain cell clusters, and 2) differentially down-regulated genes, FDR-adjusted p-value < 0.05 and log2fold-change > -0.25 throughout all cell clusters in the tau P251L *Drosophila* brain. GeneRIF terms were used for gene ontology analysis. We showed the top 5 enriched terms for all the gene enrichment analysis and removed the redundant terms in the same gene sets. Enrichment analysis data is presented as a combined score, which is the combination of the p-value (computed using Fisher exact test) and z-score (computed to assess the deviation from the expected rank) calculated by multiplying the two scores, giving the combined score (c) = log(p)*z (Chen et al. 2013). Gene enrichment terms, genes, genes contributing to the identified enrichment terms, p-values, z-scores, and combined scores are given in each supplementary files and displayed in the corresponding figures.

**Protein-protein interaction network and cell-cell communication analysis**

For the protein-protein network analysis, all the annotated clusters in the single-cell RNA sequencing dataset were classified based on their location within the *Drosophila* brain: central body, optic lobe, and glial cells. Differentially expressed genes with a log2 fold change > 0.25 and < -0.25 with FDR < 0.05, were used as input for the OmicsIntegrator protein-protein interaction package (Tuncbag et al. 2016). Fold change was used as the prize. Network hyperparameters were sorted hierarchically by the average node specificity, node robustness, and KS statistic of the degree of predicted nodes to that of prize nodes to choose a parameter set. In short, robustness was determined as the percentage of times a node appeared in the network after 100 random permutations of the network edges. Node specificity was the percentage of networks in which the node was observed after randomly shuffling the prize values and re-running the algorithm 100 times. A network parameter set corresponding to smaller specificity values, higher robustness values, and smaller KS statistics was selected. The genes identified in each network were used to annotate the biological processes using FlyEnrichr (Kuleshov et al. 2016). The cell-cell interaction analysis was performed as previously published (Liu et al. 2022).

**Trajectory analysis**

Trajectory analysis was performed using Slingshot (Street et al. 2018) on astrocyte subclusters. Trajectories were calculated on defined subclusters with the cell type showing the highest entropy used as a starting cluster (Guo et al. 2017). Differential gene expression along each pseudotime lineage was calculated using a generalized additive model (Chambers 1997). The first 1000 genes from all lineages were calculated and top 100 genes were shown on the heatmaps.

**Gene regulatory network analyses**

The previously published pySCENIC (v 0.10.0) (Single-Cell rEgulatory Network Inference and Clustering) pipeline with the *Drosophila* genome 9 (Dm9) reference genome was used to assess gene regulatory networks (Van de Sande et al. 2020). The level of regulon activity in each cell was scored using AUCell, which was then converted to a binary scale to reflect the presence or absence of the regulon.
